# Landscape viromics of introduced honeybees and bumblebees reveal distinct environmental and host-specific effects

**DOI:** 10.64898/2026.03.26.714648

**Authors:** Sabrina Haque, Emily J Remnant, James E Damayo, Fleur Ponton, Rachael Y Dudaniec

## Abstract

Understanding how viral communities vary across co-occurring hosts and environments is essential for assessing species-specific viral risks under changing land use and climate. This is particularly relevant for managing introduced bees, which face persistent viral threats themselves, as well as transmitting plant viruses. Here, we compare RNA viromes of the long-established honeybee (*Apis mellifera*, introduced to Tasmania in 1831) and the more recent invader, the bumblebee (*Bombus terrestris*, invasive since 1992), across 14 Tasmanian sites – an island still free of the viral vector, *Varroa destructor*. Using a metatranscriptomic approach on total RNA from whole bees, we identified insect- and plant-associated viruses and inferred phylogenetic patterns of insect viral sharing, divergence, and potential cross-species transmission. We also assessed spatial and environmental drivers of viral composition, diversity, and richness. Geographic longitude, precipitation, temperature, and pasture percentage influenced the total, insect-, and plant-associated viromes of *B. terrestris*. In contrast, for *A. mellifera*, only precipitation and temperature were associated with insect and plant viral alpha diversity and community composition. Phylogenetic analyses revealed that Black Queen Cell virus in *A. mellifera* from Tasmania has diverged from mainland Australian sequences, and two distinct sub-strains of Lake Sinai virus 1 were shared by both bee species. Lake Sinai virus 3 showed evidence of interspecies transmission between *A. mellifera* and *B. terrestris*. Notably, this study provides the first detection of Moku virus in Australian bees and globally in bumblebees, suggesting potential interspecies transmission among social Hymenoptera. Overall, our findings demonstrate local viral diversification and reveal that *B. terrestris* viromes are more strongly shaped by environmental factors than those of *A. mellifera*, underscoring the importance of monitoring invasive pollinators as reservoirs and vectors of viral emergence.

## INTRODUCTION

Shifts in species distributions due to climate change, urbanisation and global trade are collectively contributing to increased impacts of biological invasions that threaten biodiversity (Wilson et al., 2009; Pecl et al., 2017). Alongside biological invasions is a rise in pathogen host shifts, where novel pathogens are transmitted between species, leading to increased emergence of infectious diseases (Woolhouse et al., 2005; Longdon et al., 2014; Faillace et al., 2017). Of particular note are RNA viruses, which adapt rapidly due to high mutation rates (Holmes, 2008), enabling them to infect novel hosts and enhance transmission (Cleaveland et al., 2001; Taylor et al., 2001; Davies & Pederson, 2008). Human-mediated species invasions can facilitate the spread of viruses into new environments, leading to spill over into native species (Malmstrom et al., 2005). Further, invasive hosts may evade native pathogens or serve as viral reservoirs, increasing the risk of cross-species transmission (Rúa et al., 2011). Studying viral diversification and transmission in invasive or introduced hosts is crucial for understanding the broader ecological and evolutionary processes underlying successful biological invasions.

An emerging concern is the deliberate or accidental introduction of non-native insects for biocontrol, pollination, or farming, which can unintentionally introduce novel viral pathogens that spill over into native insect communities (Manley et al., 2015) or infect plants, in the case of plant-associated viruses (Vilcinskas, 2019). Insects can act as both hosts and vectors for a wide range of RNA viruses (Wang et al., 2023), which may be transmitted vertically from parent to offspring (Patterson et al., 2020; Wan et al., 2023) or horizontally via contact or on plants (Jia et al., 2021; An et al., 2023). Insect viruses can influence host fitness via effects on development (Xu et al., 2020), immunity (Xu et al., 2014), and sex ratios (Wang et al., 2017). Insects, such as bees, planthoppers, leafhoppers, aphids, whiteflies and mosquitoes, can also act as vectors for a wide variety of arthropod-borne pathogenic viruses (arboviruses; Hogenhout et al., 2008; Wei & Li, 2016). Plant-infecting viruses that circulate in insect vectors – such as Rice Dwarf virus in the leafhopper *Nephotettix cincticeps* – can replicate within and invade insect tissues before being transmitted via the salivary glands (Chen et al., 2011). Additionally, plant viruses vectored by insects can manipulate host behaviour, as seen in Barley Yellow Dwarf virus, which alter the behaviour of its aphid vector, *Rhopalosiphum padi*, to enhance transmission (Ingwell et al., 2012). These complex virus–host–vector dynamics highlight the need for careful assessment of virus risks associated with insect introductions, particularly as such interactions may affect both agricultural productivity and ecological interactions.

Viruses are important contributors to declining honeybee health and global colony losses (Ellis & Munn, 2005; Evans & Schwarz 2011; Mondet et al., 2014), with most known honeybee viruses being positive-sense, single-stranded RNA viruses (ssRNA; Baker & Schroeder, 2008; Brutscher et al., 2016). Moreover, bees can serve as vectors of plant viruses, the majority of which are also ssRNA viruses (Islam et al., 2020). The European honeybee (*Apis mellifera*) is a well-established reservoir of RNA viruses, many of which can spill over into wild non-*Apis* species (Tehel et al., 2016) including bumblebees, solitary bees, hoverflies, wasps, and ants (Evison et al., 2012; Fürst et al., 2014; Ravoet et al., 2014; McMahon et al., 2015). The prevalence and pathogenicity of honeybee viruses, particularly those belonging to the order of *Picornavirales* (Chen & Siede, 2007), are strongly influenced by the ectoparasitic mite, *Varroa destructor* (hereafter, *Varroa*; Martin et al., 2012; Mordecai et al., 2015; Wilfert et al., 2016), a major biological vector that feeds on honeybee fat body tissues (Ramsey et al., 2018) and has significantly altered viral dynamics in honeybee populations (Genersch & Aubert, 2010; Ryabov et al., 2014). The global spread of *Varroa*, largely driven by human-mediated movement of honeybee colonies, has facilitated the co-dispersal of highly virulent viruses, contributing to widespread declines of both managed and wild *A. mellifera* populations, as well as other pollinators (Martin et al., 2012; Wilfert et al., 2016; Rosenkranz et al., 2020). As such, investigating host-virus dynamics in introduced honeybee populations and the potential for interspecies viral spill over is critical for understanding and mitigating emerging ecological and biosecurity threats.

Most plant viruses carried by bees do not replicate within their insect hosts but are instead mechanically vectored, which means they can be transferred passively during pollen collection and floral visits (Kolliopoulou et al., 2020; Fetters et al., 2022). However, some plant viruses can infect and replicate within bees. A notable example is Tobacco Ringspot virus (TRSV), an RNA virus traditionally transmitted by nematodes and insects, such as aphids and thrips, but also shown to infect *A. mellifera* (Li et al., 2014; Flenniken, 2014). TRSV was detected throughout the body of the honeybee and was more prevalent in weaker colonies, suggesting a potential role in honeybee colony health decline (Li et al., 2014). Although there is a lack in conclusive evidence for replication of plant viruses in honeybees (Miller et al., 2014), there remains unanswered questions regarding the possible impact of plant viruses on bee physiology, immunity and interactions with other stressors. Despite these insights, the broader ecological consequences of plant virus– pollinator interactions remain unclear, underscoring the need for further research into their effects on pollinator health in changing landscapes.

Virus transmission in bees, commonly occurring through the faecal-oral route at shared floral resources (Yañez et al., 2020), is influenced by environmental factors such as temperature, UV exposure (Alger et al., 2019; Figueroa et al., 2019), and precipitation which affect virus persistence on floral surfaces (McArt et al., 2014). By altering vegetation phenology (Zhang et al., 2007), floral diversity (Suggitt et al., 2019), and resource quality (Rering et al., 2020; Descamps et al., 2021), climate change can also reshape pollinator foraging behaviour as well as viral transmission networks (Daughenbaugh et al., 2021). These changes can in turn impact bee immunity and infection susceptibility (Alaux et al., 2010; Roger et al., 2017). Environmental factors are especially critical when viruses enter new ecosystems through invasive hosts (McMahon et al., 2018). Our understanding of how environmental factors drive virus emergence remains limited yet is important for assessing the risks posed by introduced pollinators and their emerging viruses to pollination networks under variable landscape and climatic conditions.

*A. mellifera*, introduced to mainland Australia in the 1820s for honey production and crop pollination, is now a widespread generalist pollinator found across all Australian states and territories (Goulson, 2003; Prendergast et al., 2021). Managed colonies range from small-scale beekeeping to large commercial operations. In addition, feral and unmanaged colonies persist in areas with moderate climates and suitable habitats, where *A. mellifera* can is considered invasive (Moritz et al., 2005). In contrast, the European buff-tailed bumblebee (*Bombus terrestris*) was first recorded in the island state of Tasmania in 1992 and quickly established itself as an invasive species, although it remains absent from mainland Australia (Semmens et al., 1993; Hingston, 2006). Until recently, Australia was one of the last major regions free from *Varroa*, but the mite was detected on the mainland east coast in June 2022 (Chapman et al., 2023). The island state of Tasmania hitherto remains *Varroa*-free and has adopted a strategic plan to maintain this status (Tasmanian Government – Department of Natural Resources and Environment, 2025). Although *Varroa* does not parasitise bumblebees, it can facilitate viral spill over from infected honeybees to co-foraging bumblebees (Carvell, 2002; Genersch et al., 2006; Iwasaki et al., 2015). Hence, an understanding of viral dynamics and sharing in these introduced and ubiquitous bee species in Tasmania can offer a rare and time-sensitive snapshot of a pre-*Varroa* viral landscape. Simultaneously, this system can reveal the potential for viral spill over to or from the recently invasive and highly abundant *B. terrestris*, which may signify further risks to Australia’s native insect pollinators in the present or upon *Varroa’s* imminent arrival.

Here we investigate the diversity, spill over, and evolutionary relationships of RNA viruses within the long-established *A. mellifera* (introduced to Tasmania in 1831; Oldroyd et al., 1995) and the more recently invasive *B. terrestris* (introduced in 1992), sampled across 14 locations spanning variable environments across Tasmania. Using a landscape-scale metatranscriptomic approach on total RNA extracted from whole bees, we addressed two key questions: (i) Do local environmental variables influence the diversity and composition of insect- and plant-associated viruses, and does this differ between species? (ii) How do patterns of evolutionary divergence differ for insect viruses across the landscape and between species, and is there evidence for cross-species transmission? Our study uncovers ecological and evolutionary patterns of viral sharing between co-occurring introduced pollinators and the role of environment in shaping pollinator viromes, prior to the anticipated arrival of *Varroa* within the novel landscape of Tasmania. Further, by examining the unique patterns of insect- and plant-associated RNA viruses separately, we disentangle the host-specific contributions of each viral group to the total bee virome and how they differ in diversity, viral sharing, and environmental response.

## MATERIALS AND METHODS

### Study design and bee sampling

Female worker honeybees (*A. mellifera*) and bumblebees (*B. terrestris)* were collected from a total of 14 sites across Tasmania, Australia, during the active summer flight period, from January 24^th^ to February 1^st^, 2023 (Fig. 1, Table 1). *A. mellifera* workers were caught from all 14 sites while *B. terrestris* was sampled from 13 sites (Fig. 1, Table 1). The sites were selected based on prior records of the species’ presence (Hingston et al., 2002; Hingston, 2006), locations from recent previous studies (Kardum Hjort et al., 2023, 2024), and opportunistic sightings. Bees were caught within open areas of residential, urban, and rural regions with flowering plants, including road verges, grassy patches with flowers, coastal meadows, forest edges, national parks, and public gardens (as in Chapter 3). At least eight to 10 worker bees from each species were collected per site into screw-capped jars with drilled ventilation holes. Sampling at each site was conducted up to 90 minutes, with a maximum of 10 bees per species captured. Once collected, the bees were stored in a car refrigerator (∼4°C) to induce chill coma during transport and then euthanised in a freezer (–18°C) for 1–2 hours. To confirm the sex of our *B. terrestris* samples, the number of segments on the flagellum of the antennae of each sample was counted under a dissecting microscope (males have 11 segments and females have 10 segments after the pedicel). Only female worker bees were retained in the study. Bees were then chopped into small pieces and preserved in RNA Later solution at ∼4 for approximately 2-13 days, *until* returning to the laboratory where they were transferred to a –80 freezer.

**Figure 1.**
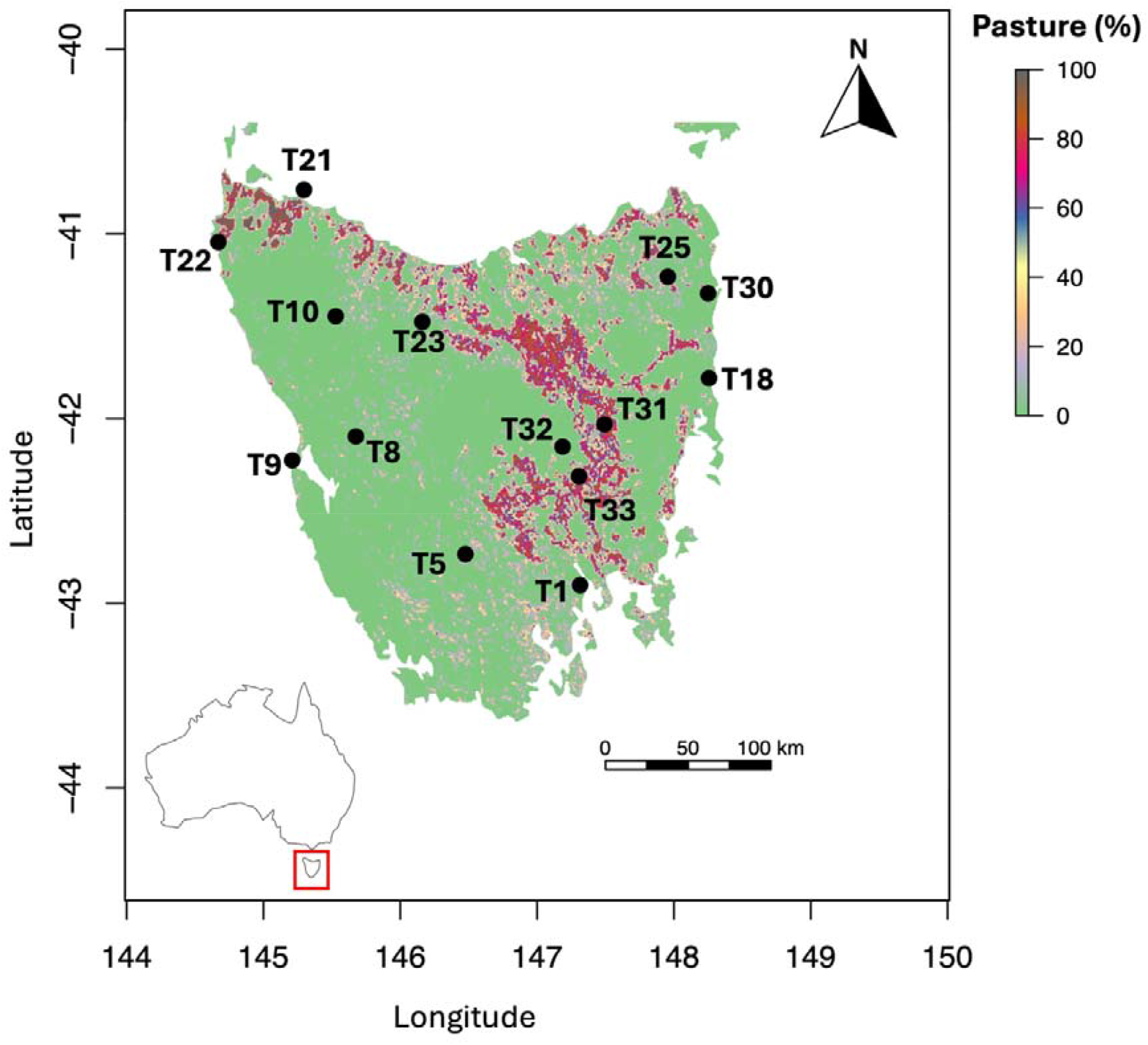
Sampling sites for *A. mellifera* (N=14) and *B. terrestris* (N=13) shown across Tasmania, Australia (inset). The sites are overlaid on a map of percentage of pasture (%). *A. mellifera* was sampled from all sites while *B. terrestris* was collected from all sites except site T18. Refer to Table 1 for corresponding site names and environmental factors and refer to Table S1 for site geographic coordinates.

**Table 1.** Site and environmental data for *A. mellifera* and *B. terrestris* across Tasmania. *A. mellifera* workers were sampled from all 14 sites while those of *B. terrestris* were collected from all sites except site T18. Abbreviations: N Am = Number of *A. mellifera* individuals pooled per site for RNA sequencing, N Bt = Number of *B. terrestris* individuals pooled per site for RNA sequencing, Temp = Mean annual temperature (C), Rain = Mean annual precipitation (mm), Wind = Average summer wind velocity (m/s), Pasture = Percentage of pasture (%), N/A = Not applicable. Refer to Fig. 1 for corresponding site locations.

| Site | Site name | N Am | N Bt | Temp | Rain | Wind | Pasture |
| --- | --- | --- | --- | --- | --- | --- | --- |
| T1 | Hobart | 10 | 4 | 11.48 | 752 | 5.8 | 0 |
| T5 | South West | 7 | 8 | 8.88 | 1632 | 5.1 | 6.06 |
| T8 | Franklin Gordon | 3 | 10 | 10.63 | 2605 | 5.6 | 0 |
| T9 | Macquarie Heads | 8 | 10 | 11.74 | 1571 | 6.7 | 2.85 |
| T10 | Tikkawoppa Waratah | 10 | 9 | 8.88 | 2046 | 4.9 | 4.84 |
| T18 | Douglas Apley | 9 | N/A | 12.66 | 687 | 5.1 | 0 |
| T21 | Stanley | 6 | 9 | 13.03 | 1038 | 6.4 | 48.8 |
| T22 | Arthur River | 10 | 7 | 12.59 | 1166 | 6.7 | 0 |
| T23 | Cethana | 10 | 10 | 9.82 | 1478 | 4.6 | 5.08 |
| T25 | Weldborough | 8 | 9 | 9.93 | 1177 | 4.3 | 11.5 |
| T30 | St Helens | 7 | 6 | 13.15 | 766 | 5.3 | 30 |
| T31 | Ross | 4 | 7 | 11.86 | 494 | 3.5 | 76.56 |
| T32 | Interlaken | 10 | 4 | 7.96 | 804 | 4 | 0 |
| T33 | Oatlands | 7 | 9 | 10.64 | 506 | 3.9 | 71 |

### Total RNA extraction, quantification and sequencing

A Qiagen RNeasy Mini Kit was used to extract total RNA from 3–10 *A. mellifera* (N = 14 sites, Table 1) and 4–10 *B. terrestris* (N = 13 sites, Table 1) individuals per site. For each *A. mellifera* sample, chopped honeybee tissue was transferred from the RNA Later storage tube to a fresh 1.5⍰mL Eppendorf tube using sterile forceps. Subsequently, 600 µL of Buffer RLT and 6 µL of 2-mercaptoethanol were added. Glass and zirconium beads (2 mm, Lysing Matrix H; MP Biomedicals) were added to each tube and the mixture was homogenized using a TissueLyser II (Qiagen) for 4 minutes at 25 Hz. The homogenate was then left at room temperature for 10 minutes to allow bubbles to dissipate before centrifugation at 12,000 rpm for 10 minutes at 4°C. The resulting supernatant was carefully transferred to a new 2 mL Eppendorf tube, and an equal volume of 70% ethanol was added and mixed by gentle vortexing. All subsequent steps were performed following the manufacturer’s protocol.

For the larger *B. terrestris* workers, sterile forceps were used to transfer chopped tissues from the RNA Later storage tube into two separate 15 mL Falcon tubes. As we wanted to extract whole body RNA from both species, each bumblebee was divided into two portions due to its size, with each portion processed individually for RNA extraction. To each tube, 2.5 mL of Buffer RLT and 25 µL of 2-mercaptoethanol were added. The contents in both tubes were homogenized using a TissueRuptor (Qiagen) for 40–50 seconds. The tubes were then centrifuged at 4,500 rpm for 20 minutes at 4°C. Approximately 650–700 µL of the resulting supernatant from each tube was carefully transferred into two new 2 mL Eppendorf tubes and equal volumes of 70% ethanol was added and mixed gently by vortexing. All subsequent steps were carried out following the manufacturer’s instructions. This procedure was repeated for all bumblebee samples. A negative control was also extracted for sequencing, in which all reagents were used but no bee tissue was added.

RNA quantification was performed using the Qubit RNA Broad Range (BR) Assay Kit (Invitrogen) and the Qubit 4 Fluorometer (Thermo Fisher Scientific), following the manufacturer’s guide. For both species, samples from each site (n = 3–10 *A. mellifera* workers from 14 sites; n = 4–10 *B. terrestris* workers from 13 sites Table 1) were pooled to achieve final concentrations of 100 ng/µL for *A. mellifera* and 50 ng/µL for *B. terrestris*, prior to sending them off for sequencing. At the Australian Genome Research Facility (AGRF) (Melbourne, Australia), the pooled samples were prepared into libraries with a ribosomal depletion step using the Illumina Ribo Zero Plus™ Microbiome Depletion Kit, and sequenced as 150 bp paired-end reads on an Illumina NovaSeq 6000 platform.

### RNA sequence filtering and contig assembly

Sequencing adaptors and low-quality reads were removed using TrimGalore version 0.6.10 (Krueger, 2021), and read quality was assessed with FastQC version 0.11.7 (Andrews, 2010). To quantify and remove ribosomal RNA (rRNA) reads from the dataset, trimmed reads were first mapped to a customised rRNA sequence library generated for each of *A. mellifera* and *B. terrestris* using Bowtie2 version 2.2.5 (Langmead & Salzberg, 2012), and the unmapped reads were extracted for subsequent genome alignment. For each alignment, unmapped reads were saved in a separate FASTQ file, which was then used for subsequent alignments. Unmapped reads were mapped to host reference genomes from NCBI using STAR version 2.7.8a (Amel HAv3.1, GCF_003254395.2 for *A. mellifera* and BomTerr2.1, GCF_910591885 for *B. terrestris*; Dobin et al., 2013), with the remaining unmapped reads retained for virus analysis. A *de novo* assembly of unmapped reads for each sample was conducted using Megahit version 1.1.3 (Li et al., 2015). The assembled contigs were then analysed for viral homology to known viruses with a BLASTn search against the nucleotide database (downloaded in October 2024) with BLAST+ version 2.12.0 (Camacho et al., 2009). For contigs that lacked homology to sequences within the nucleotide database, a BLASTx search was conducted with DIAMOND version 2.0.9 (Buchfink et al., 2015) against the non-redundant viral protein database (downloaded in October 2024).

### Consensus generation and virus identification

Due to insufficient strain-level resolution in the *de novo* assembled viral genomes and divergence from GenBank reference sequences, sample-specific consensus sequences were generated. These were produced by integrating viral contig data with iterative Bowtie2 alignments of reads to the corresponding reference genomes. Initially, the contigs were aligned to a reference viral genome obtained from NCBI using the ‘Map to Reference’ function in Geneious Prime version 2025.0.3 (https://www.geneious.com/), and viral consensus sequences were created for each sample using the ‘Generate Consensus Sequence’ function (Geneious Prime). For this step, the consensus caller threshold was set to 0% ‘majority,’ which allowed the incorporation of any sample-specific variants while minimising nucleotide ambiguities; if there was no coverage, the reference sequence was used. Each viral sequence in the consensus reference library was verified using BLASTn in NCBI to confirm its identity. When the top BLASTn hit showed a percentage identity of ≥90%, the virus identity was confirmed, and its accession number was recorded. Sequences with <90% identity were considered likely artifacts. These results were further cross-checked in Geneious Prime. In the following iterations, unmapped reads from each sample were aligned to their respective viral consensus reference sequences using Bowtie2, and new consensus sequences were generated using the same method. This process of consensus generation and re-alignment was repeated twice to confirm that the mapping to the consensus sequences had stabilised. Finally, after the last alignment, the number of reads mapping to each virus in each sample was extracted for further analyses as a measure of viral load with SAMtools version 1.5 (Li et al., 2009). Due to ecological and functional importance of RNA viruses in both insect and plant hosts, viral hits corresponding to bacteriophages and other DNA viruses, like *Apis mellifera* Filamentous virus, were excluded from all analyses, as accurate quantification of DNA virus levels using RNA sequencing data is not feasible. Hence, only RNA virus hits were retained to ensure that the dataset accurately reflected the composition, diversity and taxa of the RNA virome relevant to host-virus ecological interactions.

### Viral abundance estimation

To account for differences in sequencing depth and viral genome length, viral abundances were normalized using Reads per Kilobase Million (RPKM), calculated using the formula: (number of paired reads mapped to a virus within a site × 10^9^) / (total non-rRNA read pairs per site x viral contig length in kilobases). For each site, RPKM values of individual viruses were summed to obtain the total RPKM of RNA viruses per site. This normalization ensured that viral abundance estimates reflected true biological variation rather than biases introduced by uneven sequencing coverage or contig size. Read counts assigned to each virus were then used to calculate relative abundances for two types of analyses: (i) All viral hits were used to visualise the relative abundance of virus types (that is, plant- or insect-associated). Virus types were classified based on sequence homology to previously characterised viruses available in GenBank. While this homology-based approach provides a general indication of likely host association, it may not always reflect true host specificity, particularly for poorly studied or multi-host viruses. (ii) Taxonomic composition of major viruses, only those contributing more than 1% relative abundance at a site were retained and visualised. This 1% threshold was applied to reduce noise from low-abundance viruses and highlight the dominant taxa contributing to virome composition.

### Environmental variables selection and correlation

Environmental variables for each sampling site (Table 1) were obtained as outlined in Kardum Hjort et al. (2023; 2024) and Chapter 3. Data for mean annual temperature (℃), mean annual precipitation (mm), and average summer wind velocity (m/s) were sourced from WorldClim v2.1 (Fick & Hijmans, 2017). Pasture refers to land dominated by grasses and low-growing vegetation suitable for grazing livestock. The percentage of pasture at each site was gathered from the Dynamic Land Cover Dataset Version 2.1, which was extracted from a circular area with a 1 km radius around each site (Lymburner et al., 2015). These four environmental variables were chosen based on their established effects on morphology, selection, and gene flow in *B. terrestris* in Tasmania (Kardum Hjort et al., 2023; 2024), as well as their role in predicting the gut microbiome community composition and diversity of Tasmanian *A. mellifera* and *B. terrestris* in our companion studies (Chapter 2 and Chapter 3). In addition to these four environmental variables, geographic X (longitude) and Y (latitude) coordinates (Table S1) for each site were obtained using Google Maps and included as predictors to account for spatial structure.

A Pearson correlation matrix was generated in R version 4.5.0 (R Core Team, 2025) to examine relationships among six environmental variables across sampling sites for *A. mellifera* and *B. terrestris* separately. In *A. mellifera*, mean annual precipitation and longitude were moderately correlated (r = –0.61; Table S2), as was average summer wind velocity with longitude (r = –0.63; Table S2). Both values remained below the commonly accepted threshold for strong collinearity (r ≥ ±0.7). Similarly, in *B. terrestris*, average summer wind velocity and the longitude showed a moderate correlation (r = –0.68; Table S3), while pasture percentage and mean annual precipitation were also moderately negatively correlated (r = –0.60; Table S3). As none of the correlations exceeded the collinearity threshold, all six environmental variables were retained for further analyses.

### Community composition and alpha diversity

To explore viral community composition across sites, we conducted non-metric multidimensional scaling (NMDS) separately for *A. mellifera* and *B. terrestris*, based on Bray–Curtis dissimilarity matrices using the *vegan* R package version 2.6-10 (Oksanen et al., 2025). NMDS ordinations were performed for three virome categories: (i) the total virome, comprising all RNA viruses detected across sites, (ii) the insect virome, consisting of RNA viruses associated with insect hosts, and (iii) the plant virome, including RNA viruses linked to plant hosts. Environmental vectors (i.e., mean annual temperature, mean annual precipitation, percentage of pasture, average summer wind velocity as well as longitude and latitude) were overlaid on the NMDS ordination, to assess the influence of environment on viral community structure, using the *envfit* function in *vegan*.

Alpha diversity metrices, including Shannon’s diversity and Chao1 richness, were calculated for each of the three virome categories – the total virome, the insect virome and the plant virome – using the *phyloseq* R package version 1.51.0 (McMurdie & Holmes, 2013). To examine how environmental factors influence virome diversity, we tested the correlations between each alpha diversity index and the six environmental variables using linear models (*lm*) in R. The resulting relationships were visualised using the *ggplot2* R package version 3.5.2 (Wickham, 2016). We compared alpha diversity between host species by conducting paired t-tests on Shannon’s diversity and Chao1 richness values derived from the 13 shared locations where both *A. mellifera* and *B. terrestris* were sampled. This allowed a direct assessment of species-level differences in virome diversity and richness under shared environmental conditions.

### Isolation by distance analysis

To test whether sites that are geographically further apart exhibit greater dissimilarity in virome diversity and richness we performed tests of isolation by distance (IBD) using Mantel tests, implemented in the *vegan* R package. Separate Mantel tests were conducted for each bee species, and each for virome diversity (Shannon’s) and richness (Chao1) measures, with Bray–Curtis dissimilarity matrices used to calculate pairwise distances separately for the total virome, and for the insect and plant virome components. Geographic distance (m) between sites was calculated and used as the predictor variable.

### Phylogenetic tree construction

For phylogenetic analysis, we selected four RNA viruses commonly found in honeybees, bumblebees, and other social Hymenoptera (Remnant et al., 2017; Highfield et al., 2020): Black Queen Cell virus (BQCV), Sacbrood virus (SBV), Lake Sinai virus (LSV) and Moku virus. For LSV, two divergent strains (LSV-1 and LSV-3; percent similarity = 75.8%) were analysed separately. These viruses were the main insect-associated (bee) RNA viruses present in our dataset and are known to play significant roles in pollinator health. As this study represents the first report of Moku virus in Australia and the first detection in bumblebees worldwide, we conducted a protein alignment to confirm its identity before generating the phylogenetic tree. Alignment of our Moku consensus sequence with a reference isolate from NCBI (KU654789) yielded a pairwise identity of 99.2%, confirming it as true Moku virus. To generate phylogenetic trees, homologous viral genomes were retrieved from the NCBI database using BLASTn. Sequence alignments were performed using MUSCLE in Geneious Prime and manually trimmed for gaps and non-conserved regions in Geneious Prime. Maximum likelihood phylogenies for each viral species were estimated in IQ-TREE (Trifinopoulos et al., 2016) (http://iqtree.cibiv.univie.ac.at/) using a best-fitting substitution model selected by ModelFinder (Kalyaanamoorthy et al., 2017) according to Bayesian information criterion (BIC; Nguyen et al., 2015). Branch support was assessed with ‘Ultrafast bootstrap approximation’ with 1000 replicates (Hoang et al., 2018).

The resulting tree files were visualized in FigTree version 1.4.4 (https://tree.bio.ed.ac.uk/software/figtree/ ; Rambaut, 2018).

## RESULTS

### RNA sequencing data

Illumina sequencing of pooled *A. mellifera* samples from 14 sites produced a total of 44– 254 million paired reads per site (mean ∼174 million; Table S4), with an average of 21.25% (SE ±1.73%) non host rRNA reads retained for virome analysis (mean ∼38 million per site; Table S4). Similarly, *B. terrestris* samples from 13 sites yielded 127 million to 27 billion paired reads per site in total (mean ∼2 billion; Table S5), with 21.33% (SE ±0.40%) non-rRNA reads retained (mean ∼44 million per site; Table S5). These unaligned reads were considered potential viral sequences.

### Viral taxonomic profiles

The total abundance of RNA viruses in *A. mellifera* showed marked variation across Tasmania, ranging from as little as ∼0.44 RPKM to over 63,000 RPKM (excluding bacteriophages; Table S4, Fig. 2A). High viral loads (>10,000 RPKM) were observed at sites T1, T10, T23, T30, and T33. Intermediate viral loads (∼1000–2600 RPKM) were recorded at sites T5, T25, and T31, while the remaining sites exhibited comparatively low loads (∼0.44–408 RPKM; Table S4, Fig. 2A). These differences showed no consistent association with environmental or geographical patterns.

**Figure 2.**
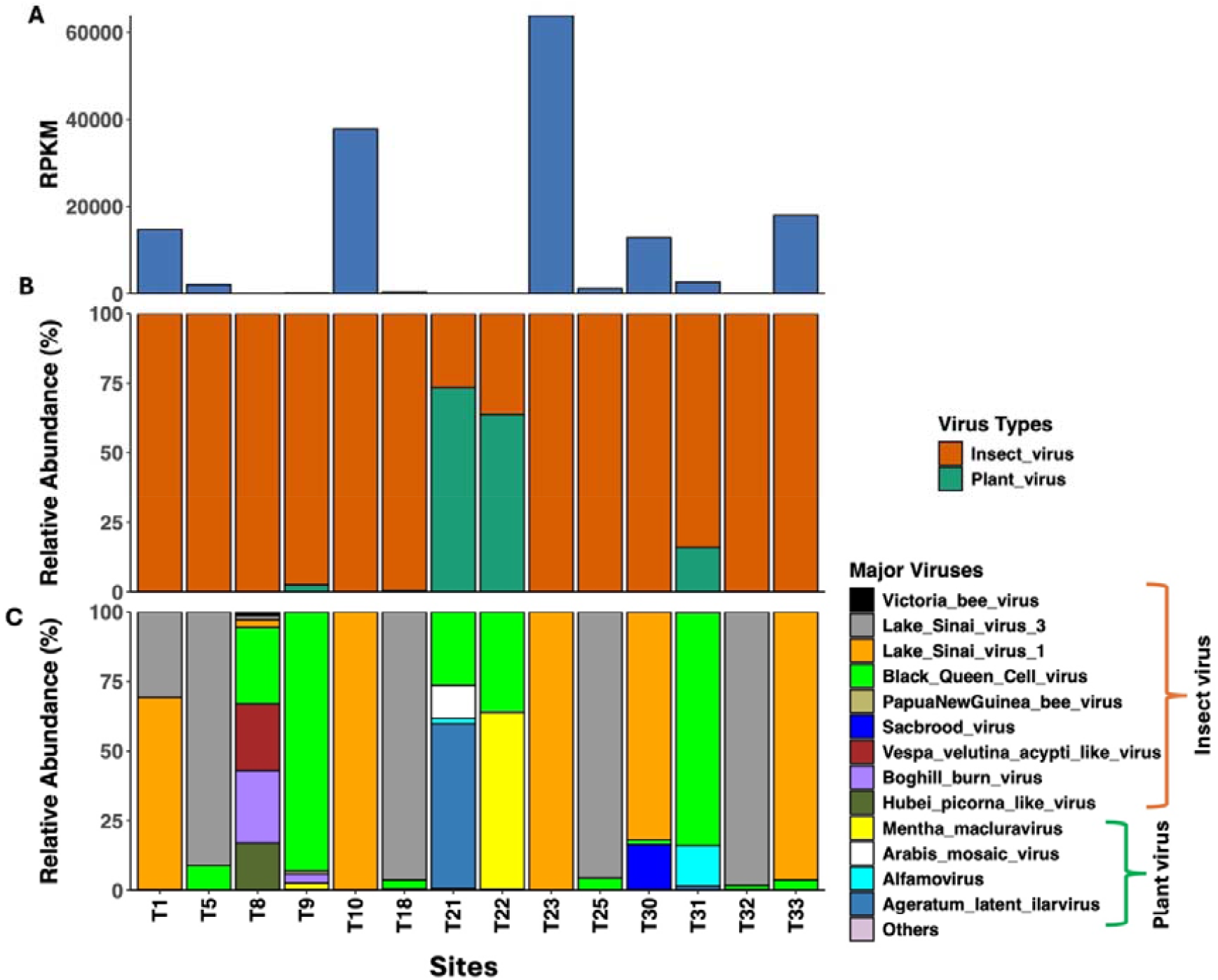
Virus abundance and composition in *A. mellifera* across Tasmania. (A) Total RNA virus abundance per site. (B) Relative abundance of virus types, showing insect and plant viruses. (C) Relative abundance of major RNA viruses (>1% at any site). In (C), the category ‘Others’ includes all RNA viruses (plant and insect) with <1% relative abundance. Study sites are shown along the x-axis in bottom panels of all three plots. Abbreviations: RPKM = Reads Per Kilobase Million.

Overall, 10 plant viral taxa and 13 insect viral taxa were identified from the virome of *A. mellifera* across all sites (Fig. S1). Insect-associated RNA viruses dominated the virome at nearly all sites, comprising ∼84–100% of the relative abundance, except for T21 and T22, where plant-associated viruses were more prominent (Fig. 2B). Taxon-level analysis revealed distinct site-specific patterns of major bee viruses in *A. mellifera* (Fig. 2C). BQCV was particularly prominent at several sites, including T5 (∼8%), T8 (∼27%), T9 (∼92%), T21 (∼26%), T22 (∼36%), and T31 (∼83%). At site T8, additional viruses such as *Vespa velutina* acypti-like virus (∼23%), Boghill Burn virus (∼26%), and Hubei picorna-like virus (∼16%) were also abundant. LSVs were widely detected across multiple sites, with LSV-1 highly prevalent in T1 (∼69%), T10 (∼99%), T23 (∼99%), T30 (∼81%), and T33 (∼96%), while LSV-3 was abundant in T1 (∼30%), T5 (∼91%), T18 (∼96%), T25 (∼95%), and T32 (∼98%). SBV was also present at T30 (∼16%). Among the plant viruses detected in *A. mellifera*, Mentha macluravirus was highly abundant at T22 (∼63%) and Ageratum Latent Ilarvirus at T21 (∼59%), with both T21 and T22 characterised by plant virus dominance. Additional plant viruses included Arabis Mosaic virus at T21 (∼11%) and Alfamovirus at T31 (∼14%).

In contrast to *A. mellifera, B. terrestris* exhibited substantially lower total RNA virus relative abundance across sites, with most locations ranging between ∼0.1 and 34 RPKM (excluding bacteriophages; Table S5, Fig. 3A). Two exceptions included site T21, which showed the highest viral load (∼111 RPKM), and T8, with the lowest (∼0.002 RPKM). Similar to *A. mellifera*, the virome of *B. terrestris* comprised 10 plant viral taxa and 13 insect viral taxa, respectively (Fig. S2). While insect-associated viruses dominated the virome at many sites (e.g., T1, T5, T8, T9, T10, T22, T23 and T30), accounting for ∼74–98% relative abundance, these sites showed no clear spatial structuring (Fig. 3B) In contrast, several locations – including T21, T25, T31, T32, and T33 were primarily composed of plant viruses (∼59–99% relative abundance), again without distinct geographic patterns (Fig. 3B).

**Figure 3.**
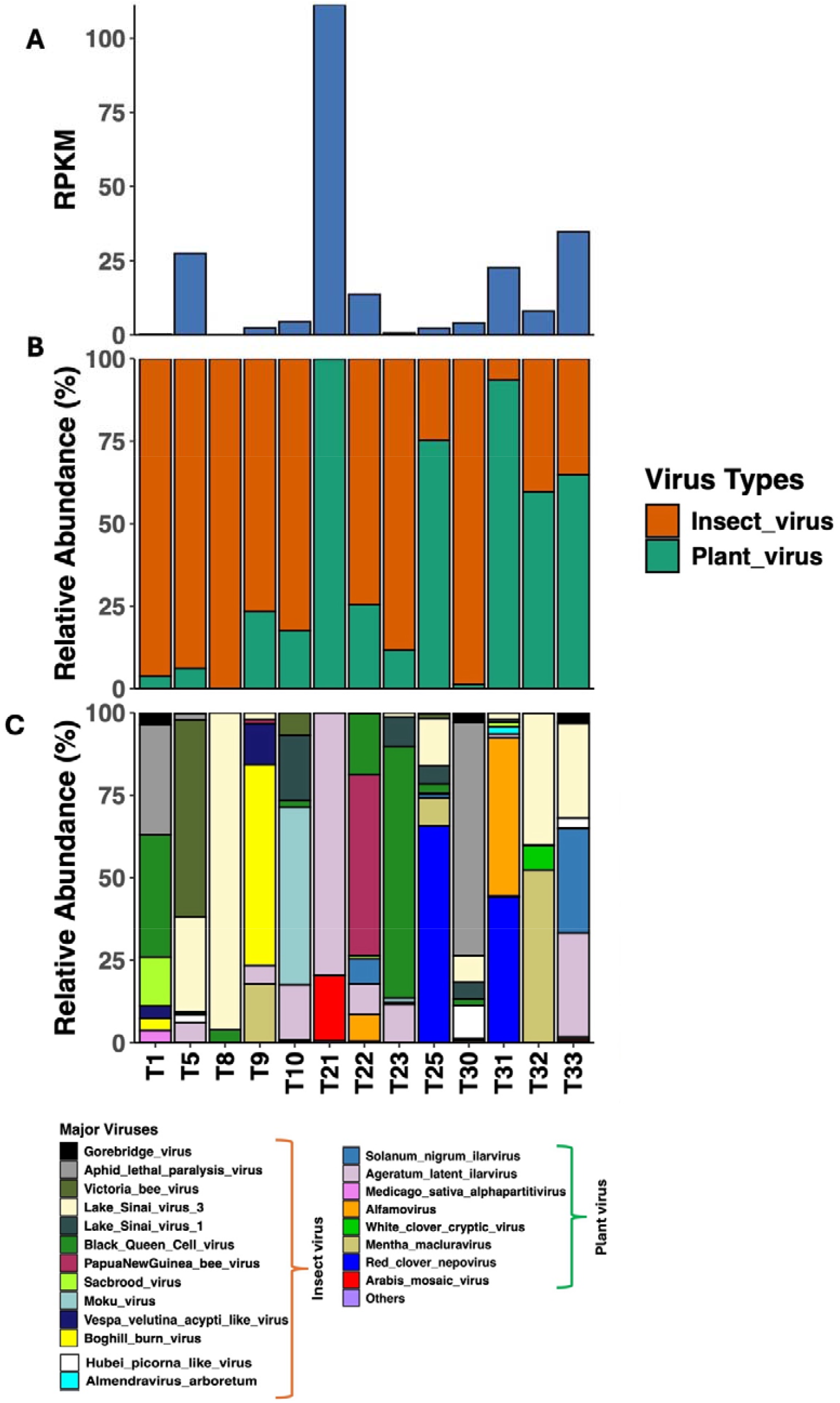
Virus abundance and composition in *B. terrestris* across Tasmania. (A) Total RNA virus abundance per site. (B) Relative abundance of virus types, showing insect and plant viruses. (C) Relative abundance of major RNA viruses (>1% at any site). In (C), the category ‘Others’ includes all RNA viruses (plant and insect) with <1% relative abundance. Study sites are shown along the x-axis in bottom panels of all three plots. Abbreviations: RPKM = Reads Per Kilobase Million.

At the taxon level, viral composition in *B. terrestris* varied markedly across sites (Fig. 3C). Key insect-associated viruses were represented at a single or few sites, including Moku virus (∼53%) and LSV-1 (∼19%) at T10; BQCV at T1 (∼37%) and T23 (∼76%); LSV-3 at T5 (∼28%), T8 (∼96%) T32 (∼39%), T33 (∼28%), T25 (∼14%) and T30 (∼8%); SBV (∼14%) at T1; Papua New Guinea Bee virus (∼54%) at T22; Boghill Burn virus (∼60%) at T9 and Aphid Lethal Paralysis virus (∼33%) at T1 (Fig. 3C). Prominent plant-associated viruses in *B. terrestris* included Arabis Mosaic virus (∼19%); Red Clover Nepovirus at T25 (∼65%) and T31 (∼44%); White Clover Cryptic virus at T32 (∼7%); Mentha Macluravirus at T9 (∼17%), T25 (∼8%) and T32 (∼52%); Alfamovirus at T22 (8%) and T31 (∼47%); *Solanum nigrum* Ilarvirus at T33 (31%) and Ageratum Latent Ilarvirus at T5 (6%), T9 (∼5%), T10 (∼16%), T21 (∼79%), T22 (∼9%), T23 (∼11%) and T33 (31%; Fig. 3C).

### Viral community composition and local environment

In *A. mellifera*, NMDS ordination of the total RNA virome revealed clear site-level differences in community composition (Fig. S3A). Environmental vector fitting indicated mean annual precipitation as a significant predictor of both the insect (p = 0.03, r^2^ = 0.52; Fig. 4A) and plant (p = 0.01, r^2^ = 0.56; Fig. S3B) virome communities (Table S7). By contrast, no environmental variable significantly accounted for variation in the total virome (all p > 0.05; Table S7; Fig. S3A), highlighting that precipitation exerts an influence on the insect- and plant-associated viromes independently, but not on the total virome composition.

**Figure 4.**
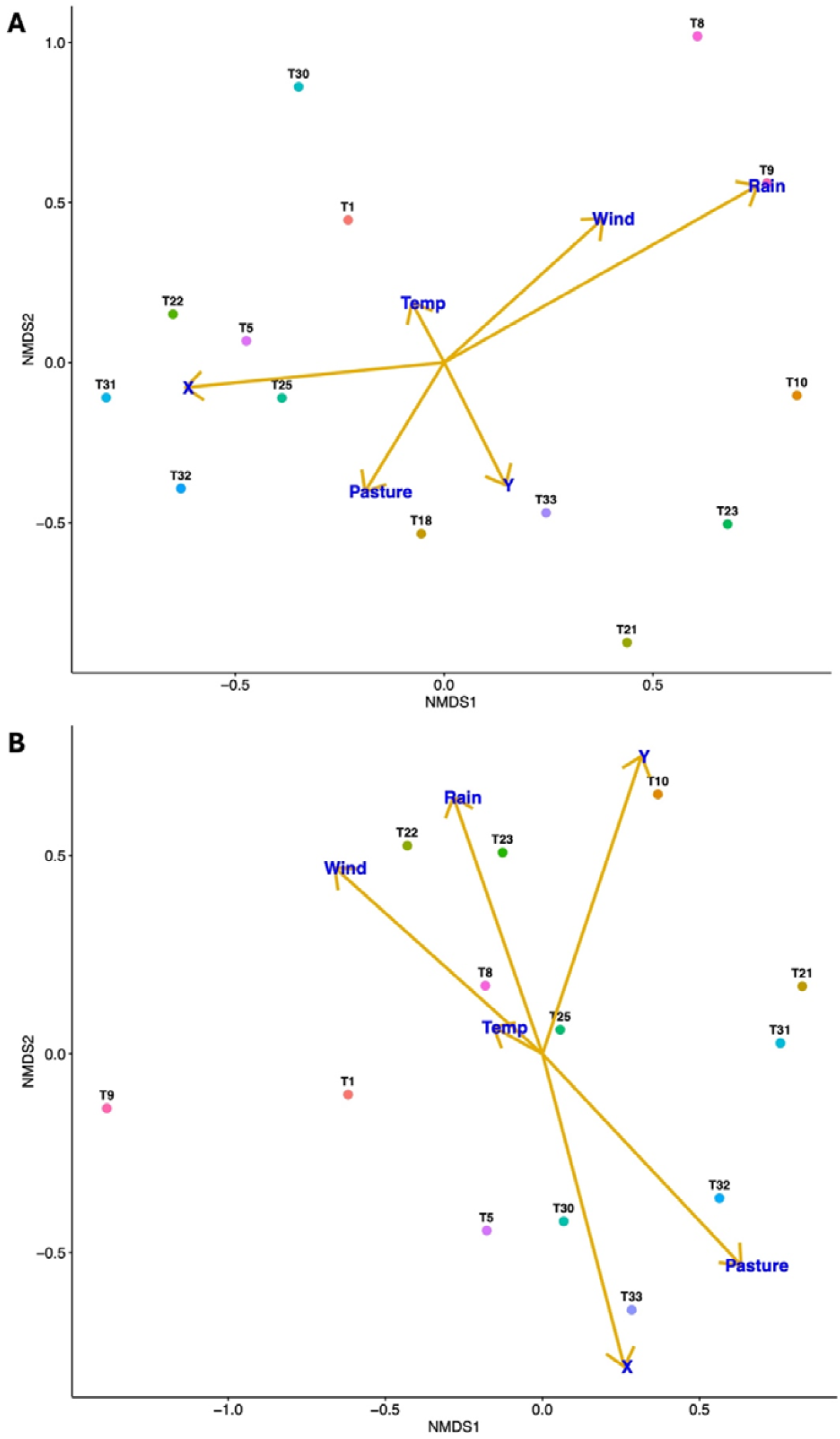
NMDS ordinations illustrating insect–viral community composition across Tasmanian sites based on Bray–Curtis dissimilarities in (A) *A. mellifera* (Stress = 0.12) and (B) *B. terrestris* (Stress = 0.16). For *A. mellifera*, mean annual precipitation (r^2^=0.52, p=0.01) was significant, whereas for *B. terrestris*, average summer wind velocity (r^2^=0.43, p=0.04), pasture percentage (r^2^=0.45, p=0.05), and longitude (r^2^=0.45, p=0.05) were significant. Abbreviations: Temp = mean annual temperature (°C), Rain = mean annual precipitation (mm), Wind = average summer wind velocity (m/s), Pasture = percentage of pasture (%), X = longitude, Y = latitude. See Fig. S3 for NMDS ordinations of total and plant-associated virome compositions for both *A. mellifera* and *B. terrestris*. Environmental vector correlations (envfit) are provided in Tables S7 and S8 for *A. mellifera* and *B. terrestris*, respectively.

For *B. terrestris*, NMDS ordination showed a broad dispersion of sites across the ordination space, reflecting greater variability in viral community composition across Tasmania (Fig. S3C). Here, environmental vector fitting identified average summer wind velocity (p = 0.04, r^2^ = 0.43), pasture percentage (p = 0.05, r^2^ = 0.45), and longitude (p = 0.05, r^2^ = 0.45) as significant drivers of insect-associated virome composition (Table S8; Fig. 4B). In contrast, neither the total nor the plant-associated viromes were significantly linked to any measured environmental variables (all p > 0.05; Table S8; Fig. S3C–D), suggesting that environmental structuring in *B. terrestris* predominantly affects the insect-associated component of the virome.

### Environmental associations with viral alpha diversity

For *A. mellifera*, Shannon’s diversity of the total and plant-associated viromes showed no significant associations with any of the environmental variables (all p > 0.05; Table S9). In contrast, insect viral diversity exhibited a positive correlation with mean annual precipitation (p = 0.05, R^2^ = 0.30; Table S9; Fig S4A). Chao1 richness patterns were slightly different: richness of the total and insect viromes did not correlate with any environmental factors (all p > 0.05; Table S9), whereas plant viral richness showed a positive relationship with mean annual temperature (p = 0.05, R^2^ = 0.28; Table S9; Fig S4B) and a negative relationship with mean annual precipitation (p = 0.001, R^2^ = 0.64; Table S9; Fig S4C). No other environmental variables showed significant correlations with insect viral richness in *A. mellifera* (all p > 0.05; Table S9).

In *B. terrestris*, mean annual precipitation was negatively correlated with total viral diversity (p = 0.05, R^2^ = 0.27; Table S10; Fig. S5A) and total viral richness (p = 0.01, R^2^ = 0.45; Table S10; Fig. S5B) while pasture percentage exhibited positive relationship with total viral richness (p = 0.01, R^2^ = 0.48; Table S10; Fig. S5C). No environmental variables significantly explained insect viral diversity (all p > 0.05; Table S10). However, insect viral richness was negatively correlated with mean annual precipitation (p = 0.03, R^2^ = 0.38; Table S10; Fig. S6A) and positively correlated with longitude (p = 0.04, R^2^ = 0.31; Table S10; Fig S6B). For plant-associated viromes, Shannon’s diversity was positively related to mean annual temperature (p = 0.05, R^2^ = 0.30; Table S10; Fig. S7A) and negatively to precipitation (p = 0.05, R^2^ = 0.28; Table S10; Fig. S7B), while Chao1 richness was positively associated with pasture percentage (p = 0.01, R^2^ = 0.48; Table S10; Fig. S7C). No other environmental variables showed significant correlations with the diversity or richness of plant-associated viromes in *B. terrestris* (all p > 0.05; Table S10).

### Comparison of viral diversity between species

The diversity of viruses (Shannon’s) for the total and insect-associated viromes were significantly higher in *B. terrestris* than *A. mellifera*. Paired t-tests revealed significant differences in both total viral diversity (p = 0.0006, t = –4.64; Fig. 5A) and insect viral diversity (p = 0.0003, t = –4.91; Fig. 5B) between the two bee species. In contrast, plant viral diversity did not differ significantly between species (p = 0.43; Fig. 5C), suggesting that interspecific differences in overall viral diversity are primarily influenced by insect-associated viruses. There were no significant differences between species for total (p = 0.16; Fig. S17A), insect (p = 0.37; Fig. S17B), or plant viral richness (Chao1; p = 0.07; Fig. S17C), indicating that despite differences in viral diversity, overall viral taxonomic richness is comparable between *A. mellifera* and *B. terrestris*.

**Figure 5.**
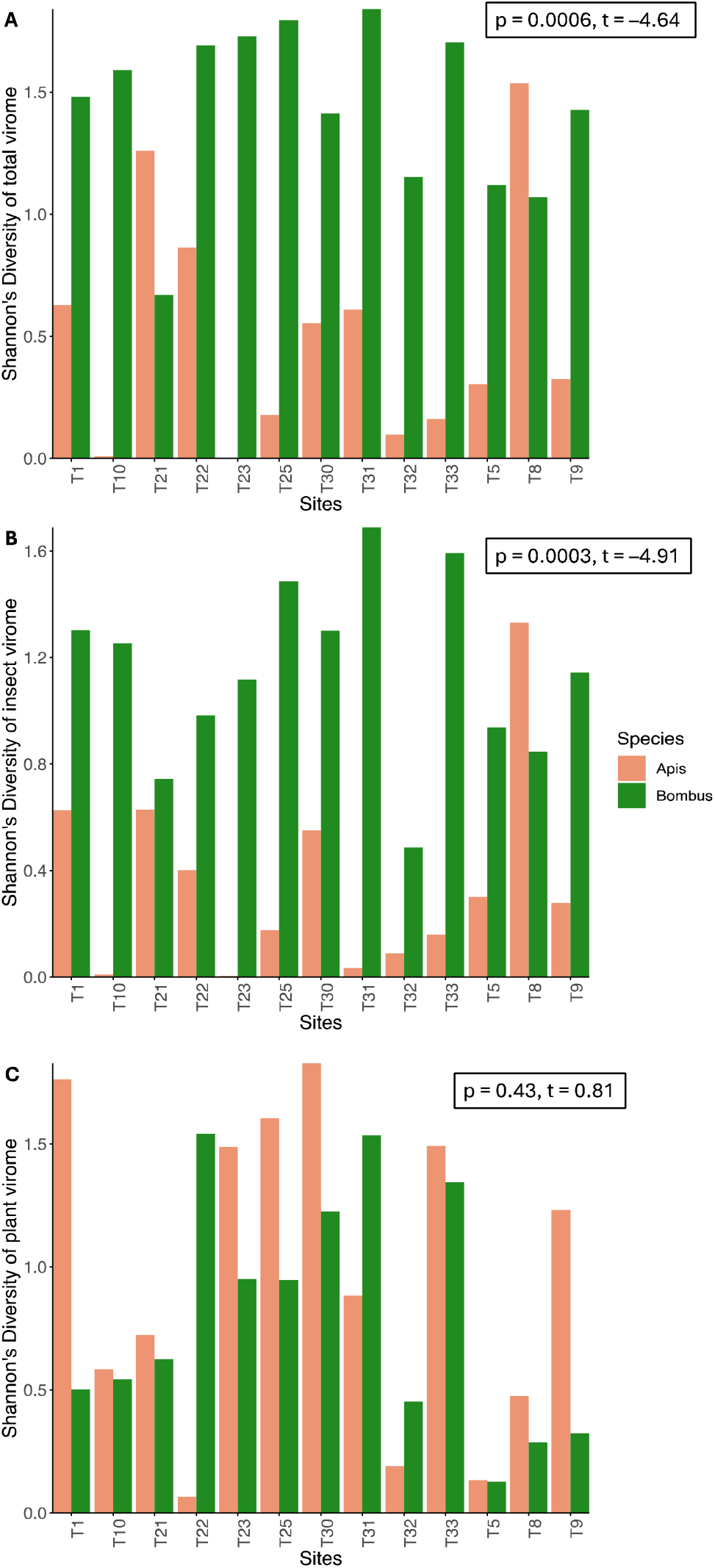
Comparison of Shannon’s diversity of viruses between *A. mellifera* and *B. terrestris* across sites: (A) the total viral diversity, (B) the insect viral diversity and (C) the plant viral diversity. Paired t-test results are shown in each panel, with significant differences observed in (A) and (B) for total and the insect virome diversity between species (p < 0.05) respectively, but not in (C) for the plant virome diversity (p > 0.05).

### Isolation by distance

There was no significant IBD relationship for total (p = 0.14), insect- (p = 0.20), or plant-associated virome diversity (Shannon’s; p = 0.66; Fig. S18) for *A. mellifera*. However, for Chao1 richness, a significant IBD relationship was found for the total virome (p = 0.01, Mantel’s r = 0.31; Fig. S19A) and plant-associated virome (p = 0.03, Mantel’s r = 0.26; Fig. S19B), but not for the insect virome (p = 0.60; Fig. S19C). In contrast, no significant IBD relationships were detected in *B. terrestris* for Shannon’s diversity: total (p = 0.52), insect (p = 0.19), or plant (p = 0.46; Fig. S20). Similarly, Chao1 richness showed no significant IBD relationship for any virome category in *B. terrestris* (all p > 0.05; Fig. S21). Therefore, virome diversity and richness in *B. terrestris* does not change with geographic distance, but in *A. mellifera* total and plant-associated virome richness do, highlighting species-specific spatial patterns in virome similarity across the sampling area.

### Phylogeny of selected bee viruses

#### Black Queen Cell virus (BQCV)

BQCV was detected in *Apis* samples from all (14) sites; however, only sequences from 12 sites were included in the phylogenetic analysis due to their higher coverage, which was sufficient for tree construction. Although BQCV was also detected in *Bombus* samples from 10 of the 13 sites, none were included in the phylogenetic tree due to insufficient sequence coverage. Phylogenetic reconstruction of BQCV sequences revealed clear biogeographic structuring, with distinct clades corresponding to Asia, Europe, and Australia (Fig. 6). Our *A. mellifera* samples, collected from Tasmania, clustered exclusively within the Australian clade, indicating that the BQCV strains in this region are evolutionarily distinct from those in the Northern Hemisphere. Within the Australian clade, our Tasmanian sequences were most closely grouped with previously published BQCV sequences from mainland Australia, including samples from Queensland, Victoria, South Australia, and Western Australia. This clustering was strongly supported by high posterior probability values, suggesting a shared evolutionary origin across the Australian continent. However, some sequence divergence was observed across Tasmania with sites T1, T23, T25, T33 and T10 forming a well-supported subclade, while others (e.g., T32, T9, T5) formed a separate Tasmanian clade (Fig. 6).

**Figure 6.**
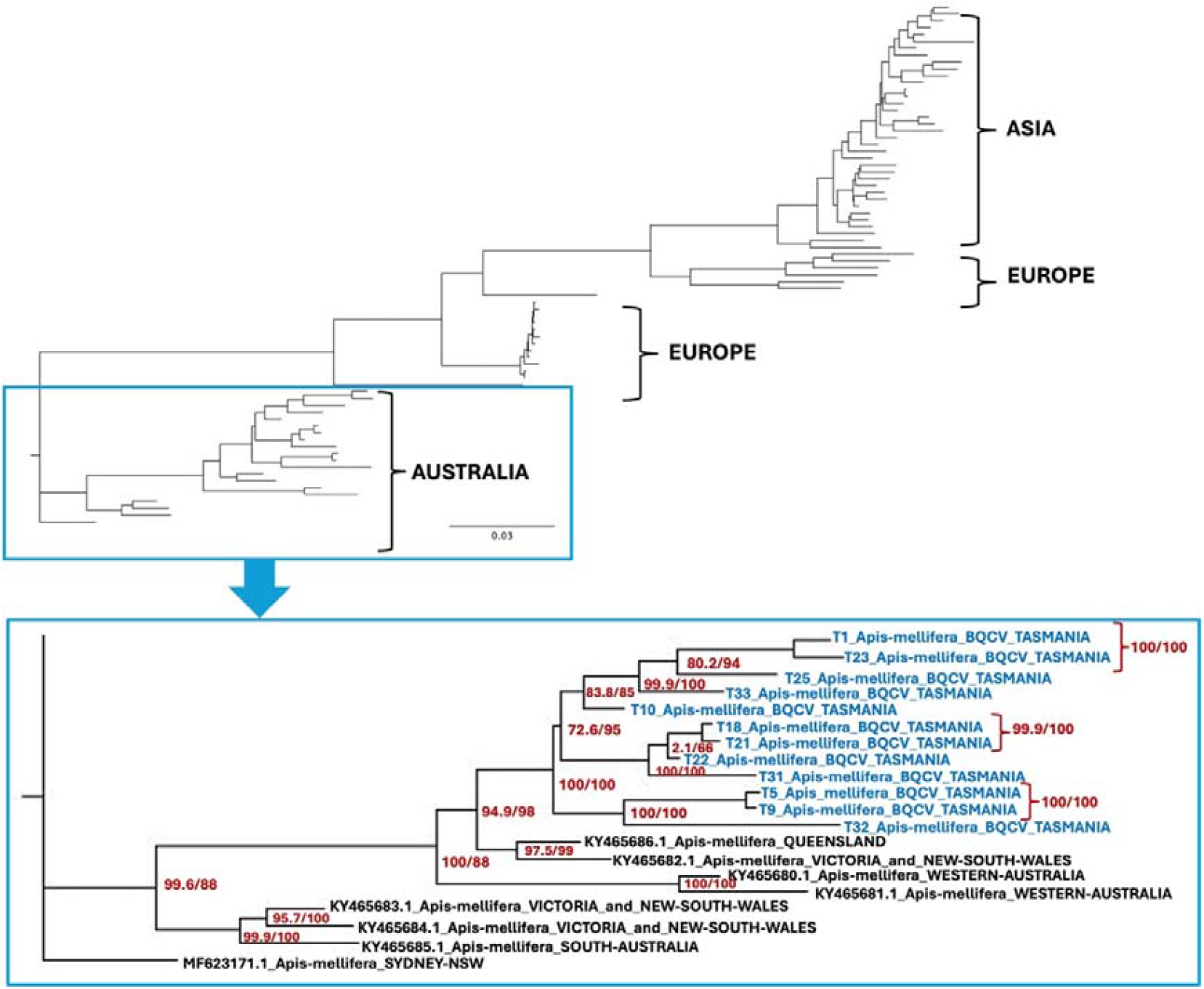
Phylogenetic tree for Black Queen Cell virus (BQCV). The phylogenetic tree was generated using maximum likelihood in IQ-TREE (Trifinopoulos et al., 2016), with the GTR+F+I+G4 substitution model which had the optimal BIC score, as determined by ModelFinder (Kalyaanamoorthy et al., 2017). Branch supports were estimated using Ultrafast bootstrap approximation (UFBoot; Hoang et al., 2017) using 1000 replicates. Support values shown are SH-like approximate likelihood ratio test (SH-aLRT) and UFBoot supports.

#### Lake Sinai virus-1 (LSV-1)

LSV-1 was found in *A. mellifera* samples from 13 of the 14 sites; however, the tree was built using sequences from six sites that had sufficient coverage for full genome construction. In *B. terrestris*, LSV-1 was present in 11 of our 13 sites, but only one sequence was included in the tree due to limited coverage. The phylogenetic tree of LSV-1, including *A. mellifera* and *B. terrestris*, showed that all Tasmanian sequences clustered within a well-supported clade alongside reference strains from the United States and Italy (Fig. 7), indicating that the LSV-1 lineage circulating in Tasmania may be a part of a globally distributed and genetically conserved viral group. Notably, a single *A. mellifera* site (T33) clustered with previously published LSV-1 sequences from mainland Australia and Tasmania, indicating phylogenetic connectivity between this Tasmanian and mainland strain which may originate from a different global source. The broader cluster of Tasmanian sequences included a *B. terrestris* sample from one site and six from *A. mellifera* from different sites that showed close sequence similarity. Notably, the *B. terrestris* sample from the north-western site T10 was most closely grouped with its geographically closest (∼50 km) *A. mellifera* sample T23, suggesting potential interspecies transmission or exposure to a common viral source. Further, *A. mellifera* samples, T1 (city of Hobart) and T30, formed well-supported subclades (Fig. 7).

**Figure 7.**
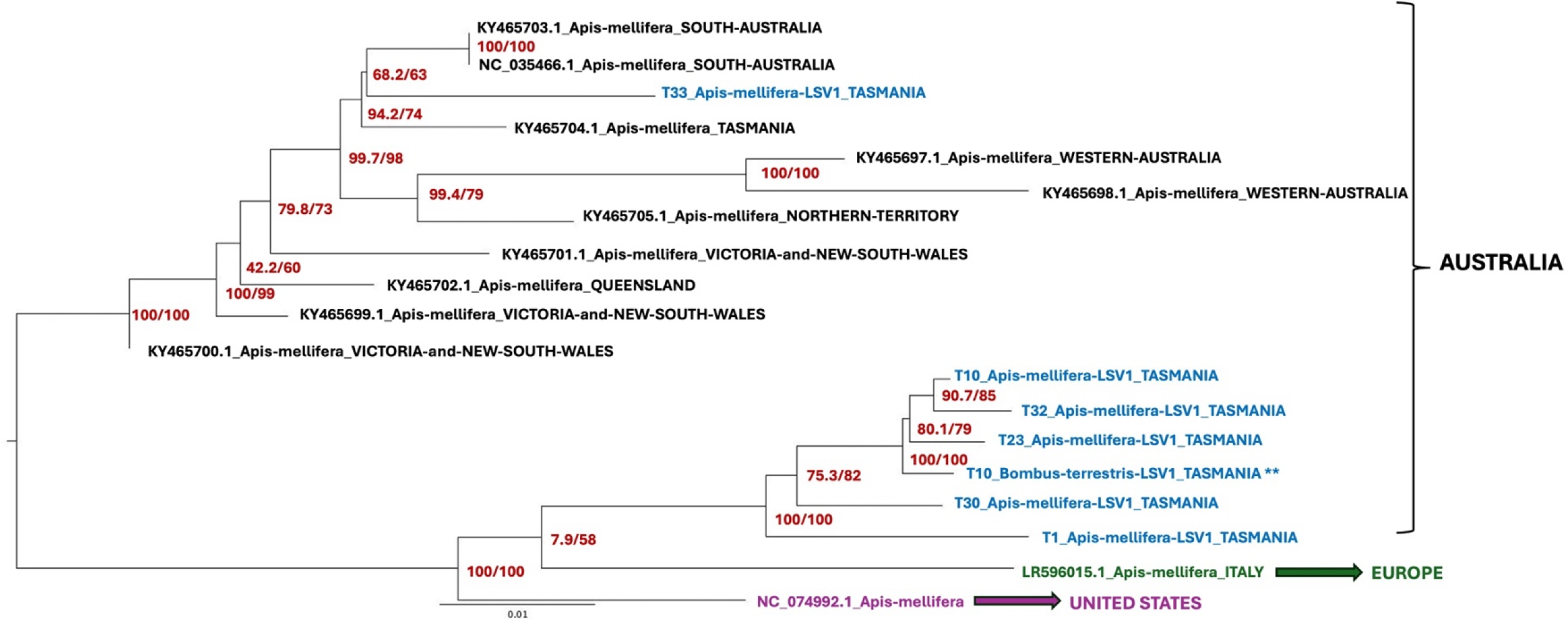
Phylogenetic tree for Lake Sinai virus-1 (LSV-1). The phylogenetic tree was generated using maximum likelihood in IQ-TREE (Trifinopoulos et al., 2016), with the TN+F+I+G4 substitution model which had the optimal BIC score, as determined by ModelFinder (Kalyaanamoorthy et al., 2017). Branch supports were estimated using Ultrafast bootstrap approximation (UFBoot; Hoang et al., 2017) using 1000 replicates. Support values shown are SH-like approximate likelihood ratio test (SH-aLRT) and UFBoot supports.

#### Lake Sinai virus-3 (LSV-3)

LSV-3 was present in *A. mellifera* samples from all (14) sites, but only nine were included in the phylogenetic analysis due to sufficient sequence coverage. In *B. terrestris*, LSV-3 was found at 10 of the 13 sites, with sequences from only two sites incorporated into the tree. Phylogenetic analysis of LSV-3 sequences showed distinct clades corresponding to samples from Europe and Asia (Fig. 8). All our Tasmanian sequences formed a well-supported monophyletic clade, clearly separated from reference sequences originating from China, Slovenia and Sweden. The two *B. terrestris* samples containing LSV-3 from T32 and T5 were grouped alongside their respective *A. mellifera* sequences from the same site, indicating minimal genetic divergence and recent, between species transmission of LSV-3 at these sites (Fig. 8). Notably, there were no reference sequences for LSV-3 available for other parts of Australia.

**Figure 8.**
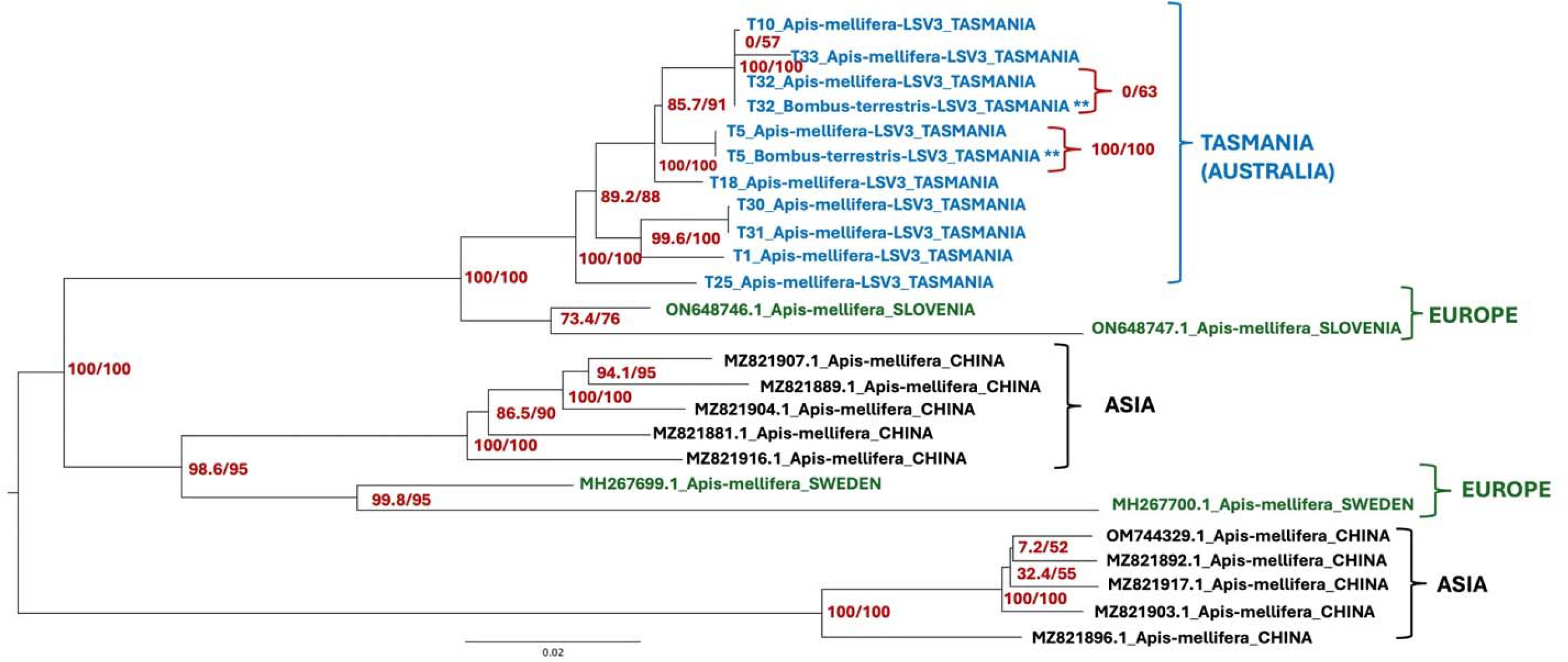
Phylogeny of Lake Sinai virus-3 (LSV-3). The phylogenetic tree was generated using maximum likelihood in IQ-TREE (Trifinopoulos et al., 2016), with the TIM3+F+G4 substitution model which had the optimal BIC score, as determined by ModelFinder (Kalyaanamoorthy et al., 2017). Branch supports were estimated using Ultrafast bootstrap approximation (UFBoot; Hoang et al., 2017) using 1000 replicates. Support values shown are SH-aLRT and UFBoot supports.

#### Sacbrood virus (SBV)

SBV was found in *A. mellifera* at 8 of the 14 sites, but only one sequence had sufficient coverage to be used in making the phylogenetic tree. In *B. terrestris*, SBV was present at 5 of the 13 sites; however, none could be used for tree construction due to insufficient coverage. The SBV phylogeny showed a clear geographic separation between Australian and Asian lineages (Fig. S22). All Australian sequences, including the SBV sample found in this study from *A. mellifera* at site T10, was placed within a well-supported and monophyletic Australian clade and was closest to a sequence previously identified in Tasmania, followed by Western Australian sequences. Other sequences in this clade were from Victoria, New South Wales, South Australia, and Western Australia, indicating that the SBV variants in Australia are geographically coherent. Reference sequences from China, Vietnam, and South Korea formed a separate and well-supported Asian clade (Fig. S22), highlighting clear phylogeographic divergence in SBV lineages across continents.

#### Moku virus

Moku virus was detected in *Apis* samples from 4 of the 14 sites, but none were included in the phylogenetic analysis due to inadequate sequence coverage. In *Bombus*, Moku virus was found at 3 of the 13 sites, with only one sequence used for generating the phylogenetic tree. The Moku virus phylogeny revealed distinct host- and geography-associated clades (Fig. 9). The Moku virus sequence from this study, obtained from *B. terrestris* in site T10, clustered within a lineage that includes sequences from the Asian hornet, *Vespa velutina*, collected in Belgium, though posterior support for the node connecting the sequences was moderate (64.9/73). This placement suggests that the Tasmanian strain is more closely related to a hornet-associated Moku virus variant than to strains detected in the Western Yellowjacket, *Vespula pensylvanica*, from Hawaii or those associated with bat metagenomes and Guangdong ant viruses from China (Fig. 9).

**Figure 9.**
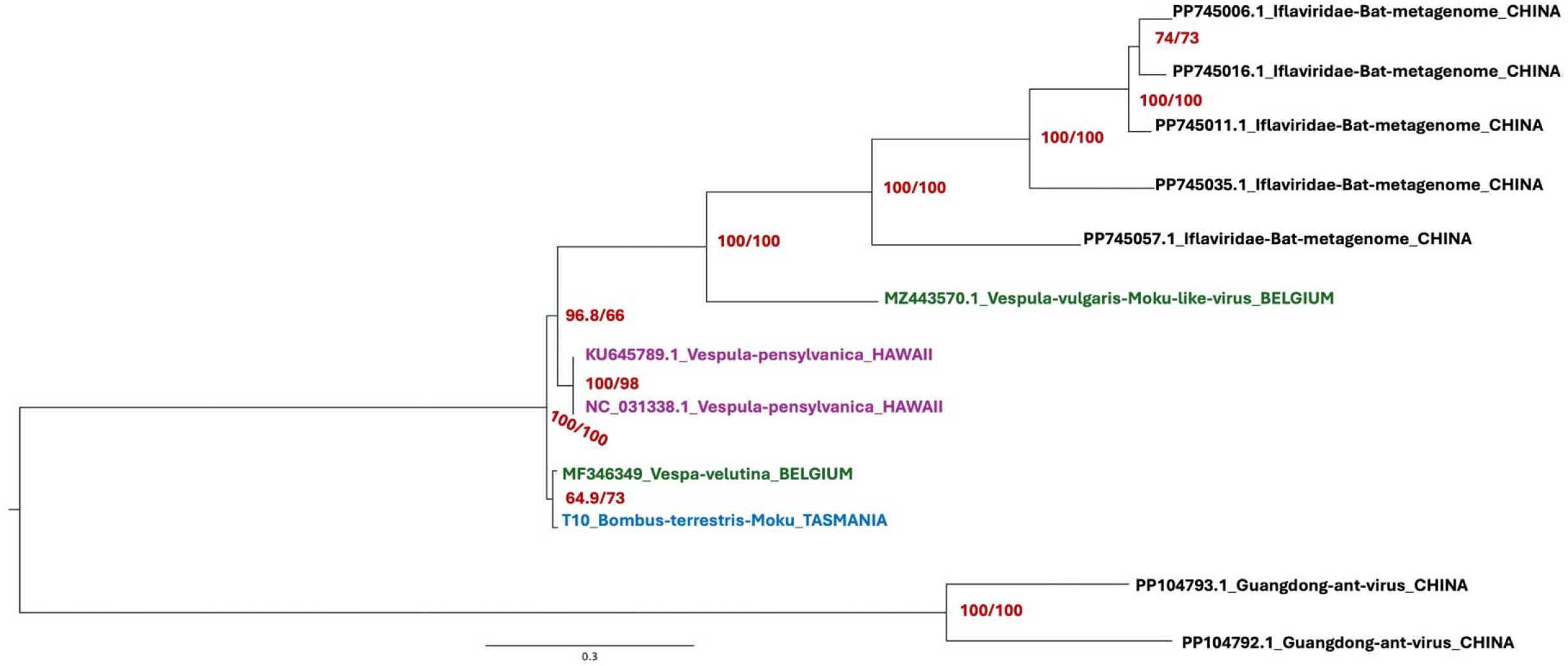
Phylogeny of Moku virus sequences. The phylogenetic tree was generated using maximum likelihood in IQ-TREE (Trifinopoulos et al., 2016), with the TVM+F+G4 substitution model which had the optimal BIC score, as determined by ModelFinder (Kalyaanamoorthy et al., 2017). Branch supports were estimated using Ultrafast bootstrap approximation (UFBoot; Hoang et al., 2017) using 1000 replicates. Support values shown are SH-like approximate likelihood ratio test (SH-aLRT) and UFBoot supports.

## DISCUSSION

Our landscape-scale virome study of *A. mellifera* and *B. terrestris* revealed clear host-specific differences in viral assemblages, shaped in part by environmental and spatial factors. In *A. mellifera*, insect (Fig. 4A) and plant-associated (Fig. S7A) viral composition were influenced by mean annual precipitation, with insect viral diversity increasing alongside precipitation (Fig. S4A) and plant viral richness increasing with temperature (Fig. S4B) but declining with higher precipitation (Fig. S4C). In *B. terrestris*, longitude, wind velocity, and pasture percentage significantly influenced insect viral composition (Fig. 4B), while precipitation was negatively correlated with total viral diversity (Fig. S5A), total viral richness (Fig. S5B), insect viral richness (Fig. S6A), and plant viral diversity (Fig. S7B). Additionally in *B. terrestris*, total viral richness increased with pasture (Fig. S5C), insect viral richness increased with longitude (Fig. S6B), plant viral diversity increased with temperature (Fig. S7A), and plant viral richness increased with pasture (Fig. S7C). Phylogenetic analysis showed that Tasmanian BQCV sequences were locally diverged (Fig. 6) and there were two distinct LSV-1 sub-strains in Tasmania (Fig. 7). Potential cross-species transmission of LSV-3 was also observed between the two bee hosts (Fig. 8). Most notably, our study marks the first ever detection of Moku virus in Australian bees and the first global report of its presence in bumblebees (Fig. 9). Altogether, our study suggests that the insect viruses are largely shaped by host identity whereas the plant viruses are more strongly influenced by the local environment.

### Effect of pasture on bee viromes

Pasture percentage was one of the key environmental drivers of insect viral community composition in *B. terrestris* (Fig. 4B). Pasture was also positively correlated to Chao1 richness of total (Fig. S5C) and plant-associated viromes (Fig. S7C) in *B. terrestris*. Bosco et al. (2024) reported that *Bombus* colonies placed in vineyard fields of varying management intensities developed more diverse viromes, with a shift from honeybee-to bumblebee-associated viruses. Colonies situated in well-connected farmland had lower viral loads, whereas those in heterogeneous, fragmented landscapes exhibited higher viral diversity. These patterns likely reflect broader anthropogenic alterations to landscape structure (Maclot et al., 2020). Agricultural intensification, urbanisation, and habitat fragmentation simplify ecosystems by reducing the availability of natural floral resources and weakening bee immunity, thereby increasing vulnerability to viral infections (Figueroa et al., 2020; Zhang et al., 2020). Pesticide and herbicide use can further reduce floral diversity and abundance (Gill et al., 2012; Harwood & Dolezal, 2020), while exposure to agrochemicals such as neonicotinoids and imidacloprid has been shown to impair immune responses and facilitate viral replication and transmission (Doublet et al., 2014; Dively et al., 2015).

Maurer et al. (2024) further demonstrated that altered landscapes with varying floral richness can disrupt bee–plant interactions critical for virus composition and transmission (Nguyen & Rehan, 2022). They found that wild bees carried higher viral loads when their floral resources overlapped with managed honeybees, while viral prevalence declined in the absence of such overlap, suggesting that flowers can act as reservoirs for viral pathogens. These dynamics may heighten the risk of cross-species transmission and the emergence of novel pathogens (Becker et al., 2015; Wallace et al., 2025), further threatening pollinator health and ecosystem stability (Beaurepaire et al., 2020). Interestingly, in our companion studies (Chapters 2 and 3), pasture percentage also significantly predicted *B. terrestris* gut bacterial composition and diversity. This highlights the role of pasture-dominated and human-modified landscapes as ecological filters that structure both microbial and viral communities. These communities may be interrelated, as gut microbiota can influence viral impacts in bees (Dosch et al., 2021), suggesting that microbiome shifts driven by modified landscapes (e.g. as found in Chapters 2 and 3) could in turn, mediate viral susceptibility and transmission. Our finding of pasture-associated effects on viral diversity therefore underscores the need to consider how interacting environmental factors shape bee health across modified landscapes.

### Local environment influences viral communities

Precipitation emerged as the most influential environmental factor for both *A. mellifera* and *B. terrestris*. In *A. mellifera*, mean annual precipitation was the primary driver of insect- (Fig. 4A) and plant-associated (Fig. S3B) viral composition, showing a positive relationship with insect viral diversity (Fig. S4A) and a negative relationship with plant viral richness (Fig. S4C). These results are consistent with prior studies showing that rainfall influences viral prevalence and transmission dynamics in bees (Piot et al., 2022). For example, high rainfall has been linked to increased prevalence of certain viruses in wild bumblebee populations (Pascall et al., 2021). Rain may also reduce faecal–oral virus transmission by limiting defecation and foraging flights, potentially concentrating viral load within colonies while decreasing inter-colony spread (McAfee et al., 2025). Reduced foraging under wet conditions can lower host mobility (Reeves et al., 2024), which may amplify local transmission (Vincze et al., 2024). Rainfall also interacts with temperature to shape viral dynamics – cooler, wetter climates limit floral resources and bee foraging (Switanek et al., 2017; Kammerer et al., 2021). Moreover, excess precipitation can dilute nectar (Mahankuda & Tiwari, 2024), disrupt pollen availability and honey production (Vincze et al., 2024), and impair suitable nesting conditions (Antoine & Forrest, 2020). Together, these findings suggest that increased rainfall, whether seasonal or driven by climate change, can disrupt bee foraging, nesting, and virus transmission.

Wind velocity was one of the significant drivers of insect viral community composition in *B. terrestris* (Fig. S4B). We previously found that wind velocity negatively correlated with core gut bacterial diversity in *B. terrestris* (Chapter 3). Wind presents physical challenges to bees due to their small size and susceptibility to turbulence (Hunter, 2007; Combes & Dudley, 2009) and has been shown to reduce foraging activity in both *A. mellifera* and *Bombus* spp. (Pinzauti, 1986; Vicens & Bosch, 2000; Hennessy et al., 2020; Hennesy et al., 2021). Reduced foraging may limit contact with virus-contaminated flowers, thus lowering inter-colony transmission, while increased defecation within the hive could elevate internal viral loads (McAfee et al., 2024; McAfee et al., 2025). Wind can also interact with temperature to exacerbate infection dynamics, such as intensifying SBV infections under high heat (McAfee et al., 2025). These findings highlight wind as a key climatic factor shaping both bee behaviour and virome characteristics.

Earlier, we found in our companion study (Chapter 3) that precipitation was also positively correlated with the diversity and richness of facultative (i.e., environmentally acquired) and total gut bacterial communities in *B. terrestris*. Precipitation was also found to interact with foraged pollen diversity to influence core and facultative gut bacterial diversity in both bee species, while wind shaped core gut bacterial richness in *terrestris*, but not *A. mellifera* (Chapter 3). These microbiome shifts, caused by environmental variables, may have downstream effects on viral communities via effects on bee immunity. Bees with intact gut microbiota are known to have higher survival rates when exposed to RNA viruses compared to those with disrupted microbiomes (Dosch et al., 2021). Core bacterial taxa such as *Snodgrassella alvi* and *Lactobacillus apis* (part of *Lactobacillus* Firm-5) can stimulate antimicrobial peptide production essential for antiviral defense (Deng et al., 2024), while environmentally acquired strains like *Apilactobacillus kunkeei* can enhance immune responses and protect against pathogenic infections (Todorov et al., 2024). Collectively, our results provide a basis for exploring the impacts of environmental factors, like precipitation and wind on bee viromes. Since the gut microbiome plays a role in viral tolerance, these findings suggest the complex ecological interplay between climate, the gut microbiome, and viral infections. While further mechanistic and experimental studies are needed, these associations offer new insight into how environmental variation may influence pollinator health at a landscape scale.

### Impact of geographical location on viral populations

We found that site longitude significantly influenced the insect viral composition (Fig. 4B) and was positively correlated with insect viral richness in *B. terrestris* (Fig. S6B), but not in *A. mellifera*. Further, total and plant viral richness in *A. mellifera* was found to increase with geographic distance between sites (Figs. S10A and S10C) – a relationship which was absent in *B. terrestris*. Previously, Manley et al., (2020) found that geographic location was a major determinant of overall viral composition and diversity in both managed *A. mellifera* and wild *B. terrestris* populations, with certain viruses exhibiting localised prevalence and others being more widespread; however, the study did not assess the role of environmental factors. Notably, Robinson et al. (2024) reported significant isolation by genetic distance in *A. mellifera* viruses (*Narnaviridae*). Shifting environmental conditions can also affect viral communities in honeybees, such as temperature and humidity, which in turn can affect viral replication and transmission dynamics (Hristov et al., 2020). For example, higher temperatures might favour the spread of certain viruses while lower temperatures could slow down viral replication in honeybees (Wang et al., 2016; Nürnberger et al., 2018). Urban and rural environments can also host distinct bee viral communities (Youngsteadt et al., 2015), as urban areas with abundant floral resources and ornamental plants support healthier bee populations and potentially more diverse viromes (Sponsler et al., 2020). In addition, these findings, along with others, highlight the importance of spatial and environmental factors in shaping bee viromes.

### Phylogenetic relationships of key bee viruses

Black Queen Cell Virus (BQCV) is one of the most widespread honeybee viruses, named for its lethal impact on queen larvae, which darken and die within their cells (Radzevičiūtė et al., 2017; Alger et al., 2019; Leat et al., 2000). Although primarily associated with *A. mellifera*, BQCV also occurs in non-*Apis* pollinators, including native Australian Hymenoptera (Brettell et al., 2020). Our phylogenetic analysis of BQCV (from *A. mellifera*) revealed local divergence within Tasmania, with sequences forming a distinct cluster from the broader Australian clade (Fig. 6). In honeybees, BQCV is transmitted via multiple pathways: horizontally through brood food, vertically from queens to offspring, through co-infection with *Nosema apis*, and potentially via drifting workers or contaminated hive materials (Allen & Ball, 1996; Chen et al., 2006; Doublet et al., 2024). While often asymptomatic in adult honeybees, high BQCV loads are frequently observed in collapsing colonies, likely due to its persistence in dead larvae, honey, and pollen for up to four weeks, which facilitates ongoing transmission (Naggar & Paxton, 2020). Studies suggest that BQCV from different geographic regions show varying degrees of genetic similarity as BQCV isolates from Japan and Thailand tend to cluster together (Mookhploy et al., 2015). Although BQCV is found across multiple *Apis* species (*A. mellifera, Apis cerana, Apis florea, Apis dorsata*), geographic origin appears to be a stronger driver of viral divergence than host species (Tapaszti et al., 2009; Mookhploy et al., 2015). Phylogenetic studies across Asia, Europe, and Australia show regional clustering of BQCV strains, reflecting both shared ancestry and location-specific evolution (Avci et al., 2022; Xu et al., 2024). Moreover, differences in transmission routes may affect the virulence and distribution of geographically distinct BQCV variants (Naggar & Paxton, 2020). In Tasmania, we observed distinct clustering of Tasmanian BQCV isolates, pointing to potential regional viral evolution which may be due to geographical isolation, environmental or host-related factors.

LSV is a genetically diverse group of honeybee viruses, consisting of LSV-1 and LSV-2 as primary strains, along with additional sister strains (such as LSV-3–8, NE, SA1, SA2, TO; Runckel et al., 2011; Nguyen et al., 2024). LSVs are highly diverse and rapidly evolving, with significant sequence variation even within the same type or strain (Cornman, 2019, Hou et al., 2023). LSVs have been detected in *Varroa* mites; however, Ravoet et al., (2015) found that LSVs do not replicate within the mites, and current evidence remains insufficient to confirm *Varroa*-mediated transmission. Their high abundance in the honeybee gut instead suggests potential faecal–oral or food-associated transmission routes (Daughenbaugh et al., 2015). Though not always linked to overt disease, LSVs are often found at higher levels in weaker colonies, suggesting a role in colony health (Cornman et al., 2012; Glenny et al., 2017). In addition, our LSV-3 phylogenetic tree revealed evidence of cross-species transmission, as *A. mellifera* and *B. terrestris* samples within two sites (T5 and T32) contained phylogenetically identical LSV-3 strains (Fig. 8). Similarly, Šimenc et al. (2020) reported LSV-3 in both *A. mellifera* and *B. terrestris*, with bumblebee-derived strains showing 98.6% to 99.4% nucleotide identity with those from honeybees, suggesting recent interspecies transmission events or a shared environmental source of infection. In constructing our LSV-3 phylogenetic tree, we found no available reference sequences from mainland Australia. The Tasmanian clade for LSV-3 grouped more closely with sequences from Slovenia (Europe) than those from Asia (Fig. 8). This current phylogeny is limited by a lack of geographically diverse reference sequences and would benefit from inclusion of mainland Australian strains, if present, to improve resolution and contextualise the position of our Tasmanian LSV-3 isolates.

This study is the first to report the presence of Moku virus in Australian bees and is the first reported case in bumblebees globally, extending both the known host range and geographic distribution of this virus. Moku virus is phylogenetically related to Slow Bee Paralysis Virus (SBPV), a highly virulent but rarely detected pathogen in honeybees and bumblebees. This relationship raises the possibility that Moku virus may have entered bumblebee populations via similar transmission pathways or mechanisms as SBPV, thus potentially sharing aspects of its pathogenic ecology (Mordecai et al., 2016). Phylogenetic analysis revealed that the Tasmanian Moku virus sequence from *B. terrestris* clustered closely with *V. velutina* (Asian hornet) sequences from Belgium (Fig. 9), suggesting potential interspecies transmission among social Hymenoptera. Such transmission is likely driven by ecological interactions, including shared floral resources and predation (Singh et al., 2010). Moku virus was originally discovered in the predatory social wasp *V. pensylvanica* (Western yellowjacket) in Hawaii, where it was detected at high abundance, suggesting that wasps may be its natural host (Mordecai et al., 2016; Highfield et al., 2020). Moku-like variants have since been identified in the common English wasp (*Vespula vulgaris*; Remnant et al., 2021) and the Asian hornet (*V. velutina*; Dalmon et al., 2019), further supporting the role of Vespids as reservoir hosts. However, frequent detections of Moku virus in *A. mellifera* (Garigliany et al., 2018; Highfield et al., 2020) and in *Varroa* mites (Mordecai et al., 2016) indicate a broad host range. Notably, Vespids such as the European wasp (*Vespula germanica*) and English wasp (*V. vulgaris*) are present in Tasmania (Department of Natural Resources and Environment Tasmania, 2025), suggesting possible regional vectors. While Moku virus has been detected in various species, its pathogenicity, particularly in wasps, honeybees, and bumblebees, remains unclear. Continued surveillance is necessary to better understand its potential impacts on pollinator health and ecosystem stability.

## CONCLUSION

We characterised and compared the RNA viromes of two co-occurring introduced pollinators, *Apis mellifera* and *Bombus terrestris*, which were introduced to Tasmania at different timepoints. Our findings show that viral community structure and evolution in novel environments are significantly shaped by host identity, climate variation, and geographic location. Future research should incorporate multi-season sampling across varying landscape and climate types to better capture temporal and spatial variation, as our study represents a single snapshot in time. Despite these limitations, this study provides a critical baseline of pollinator viromes within a Varroa-free system and underscores the need to monitor viral diversity, cross-species transmission, and environmental drivers in introduced pollinators. As climate and land-use change intensify, the likelihood of virus host shifts may increase, posing potential threats to pollinator health and ecosystem stability. Ultimately, our findings contribute to a broader understanding of how landscape and climate variation influence virus evolution and transmission in wild pollinator populations.

## Acknowledgements

We would like to thank Georgina E. Binns for fieldwork assistance and Cecilia Kardum-Hjort for prior identification of many sampling sites. This project was funded by a Macquarie University Research Acceleration Grant and an Australian Research Council Future Fellowship (FT230100478) awarded to R. Dudaniec, and graduate student funding (to S. Haque) from the School of Natural Sciences at Macquarie University. Permits for collecting *Apis mellifera* and *Bombus terrestris* were obtained from the Department of Primary Industries, Parks, Water and Environment, Tasmania (Authority No. FA22410). A permit for transportation of deceased *A. mellifera* and *B. terrestris* to New South Wales was obtained from the Department of Primary Industries, NSW Government (Ref: OUT22/16254).

