## Supplementary Materials for "Landscape viromics of introduced honeybees and bumblebees reveal distinct environmental and host-specific effects"

**SUPPLEMENTARY MATERIAL**

**Table S1.** Geographic coordinates for all study sites.

| Site ID | Site Name | Longitude  (X coordinate) | Latitude  (Y coordinate) |
| --- | --- | --- | --- |
| T1 | Hobart | 147.31423 | -42.90123 |
| T5 | South West | 146.47587 | -42.73323 |
| T8 | Franklin Gordon | 145.67533 | -42.09598 |
| T9 | Macquarie Heads | 145.2099 | -42.225 |
| T10 | Tikkawoppa Waratah | 145.52776 | -41.44534 |
| T18 | Douglas Apley | 148.25657 | -41.78077 |
| T21 | Stanley | 145.29589 | -40.76035 |
| T22 | Arthur River | 144.66993 | -41.04383 |
| T23 | Cethana | 146.16065 | -41.47527 |
| T25 | Weldborough | 147.95554 | -41.2315 |
| T30 | St Helens | 148.2498 | -41.3218 |
| T31 | Ross | 147.4922 | -42.031 |
| T32 | Interlaken | 147.18628 | -42.15115 |
| T33 | Oatlands | 147.30541 | -42.3112 |

**Table S2.** Pearson’s correlation matrix for all environmental variables in sites used for the *A. mellifera* virome study. Longitude (X) showed moderate correlations with mean annual precipitation and average summer wind velocity which are highlighted in bold. Abbreviations: Temp = Mean annual temperature ($℃)$, Rain = Mean annual precipitation (mm), Wind = Average velocity of summer wind (m/s), Pasture = Percentage of pasture (%), X = Longitude, Y = Latitude.

|  | Temp | **Rain** | **Wind** | Pasture | X |
| --- | --- | --- | --- | --- | --- |
| Temp |  |  |  |  |  |
| Rain | -0.36 |  |  |  |  |
| Wind | 0.50 | 0.33 |  |  |  |
| Pasture | 0.30 | -0.52 | -0.43 |  |  |
| **X** | 0.02 | **-0.61** | **-0.63** | 0.23 |  |
| Y | 0.36 | 0.03 | 0.21 | 0.09 | -0.23 |

**Table S3.** Pearson’s correlation matrix for all environmental variables in sites used for the *B. terrestris* virome study. Longitude (X) showed moderate correlation with average summer wind velocity while pasture percentage was moderately correlated with mean annual precipitation – both of which have been shown in bold. Abbreviations: Temp = Mean annual temperature ($℃)$, Rain = Mean annual precipitation (mm), Wind = Average velocity of summer wind (m/s), Pasture = Percentage of pasture (%), X = Longitude, Y = Latitude.

|  | Temp | **Rain** | **Wind** | Pasture | X |
| --- | --- | --- | --- | --- | --- |
| Temp |  |  |  |  |  |
| Rain | -0.31 |  |  |  |  |
| Wind | 0.53 | 0.34 |  |  |  |
| **Pasture** | 0.38 | **-0.60** | -0.44 |  |  |
| **X** | -0.10 | -0.58 | **-0.68** | 0.34 |  |
| Y | 0.37 | 0.03 | 0.21 | 0.09 | -0.26 |

**Table S4.** Summary of RNA sequencing metrics for pooled *A. mellifera* samples across sites. Reads not aligned to host rRNA represent the pool of reads potentially containing viral sequences. Abbreviations: RPKM = Reads per Kilobase Million (mapped reads), SE = standard error.

| **Sites** | **total no. of read pairs** | **% paired reads aligned to host rRNA** | **no. of read pairs aligned to host rRNA** | **% paired reads not aligned to host rRNA** | **no. of read pairs not aligned to host rRNA** | **RPKM of all RNA viruses per site**  **(excluding phages)** |
| --- | --- | --- | --- | --- | --- | --- |
| T1 | 207222377 | 59.43 | 123152259 | 40.57 | 84070118.3 | 14713.7683 |
| T5 | 169288775 | 83.15 | 140763616 | 16.85 | 28525158.6 | 2043.45255 |
| T8 | 220101370 | 84.42 | 185809577 | 15.58 | 34291793.4 | 0.4425122 |
| T9 | 184214644 | 78.10 | 143871637 | 21.90 | 40343007.0 | 143.620628 |
| T10 | 173980110 | 81.90 | 142489710 | 18.10 | 31490399.9 | 37819.4499 |
| T18 | 188120098 | 80.50 | 151436679 | 19.50 | 36683419.1 | 408.962211 |
| T21 | 232981869 | 81.60 | 190113205 | 18.40 | 42868663.9 | 48.9350227 |
| T22 | 44368968 | 79.66 | 35344319.9 | 20.34 | 9024648.09 | 37.840499 |
| T23 | 211961292 | 81.49 | 172727257 | 18.51 | 39234035.1 | 63918.1834 |
| T25 | 151277919 | 78.00 | 117996777 | 22.00 | 33281142.2 | 1128.31459 |
| T30 | 156512027 | 79.69 | 124724434 | 20.31 | 31787592.7 | 12833.6664 |
| T31 | 56975266 | 83.73 | 47705390.2 | 16.27 | 9269875.78 | 2601.08691 |
| T32 | 254160534 | 70.91 | 180225235 | 29.09 | 73935299.3 | 90.3403241 |
| T33 | 193356199 | 79.97 | 154626952 | 20.03 | 38729246.7 | 18003.2859 |
| Mean | **174608675** | **78.75**  **(SE** $\boldsymbol{\pm1.73)}$ | **136499075** | **21.25**  **(SE** $\boldsymbol{\pm1.73)}$ | **38109600** | **10985.0964** |

**Table S5.** Summary of RNA sequencing metrics for pooled *B. terrestris* samples across sites. Reads not aligned to host rRNA represent the pool of reads potentially containing viral sequences. Abbreviations: RPKM = Reads per Kilobase Million (mapped reads), SE = standard error.

| **Sites** | **total no. of read pairs** | **% paired reads aligned to host rRNA** | **no. of read pairs aligned to host rRNA** | **% paired reads not aligned to host rRNA** | **no. of read pairs not aligned to host rRNA** | **RPKM of all RNA viruses per site (excluding phages)** |
| --- | --- | --- | --- | --- | --- | --- |
| T1 | 150483240 | 76.90 | 115676466.6 | 23.13 | 34806773.4 | 0.19961063 |
| T5 | 187103336 | 79.50 | 148765862.5 | 20.49 | 38337473.5 | 27.5294429 |
| T8 | 27127315902 | 80.00 | 21704565453 | 19.99 | 5422750449 | 0.00154363 |
| T9 | 127115966 | 77.60 | 98603854.83 | 22.43 | 28512111.2 | 2.30444569 |
| T10 | 168956805 | 75.70 | 127832718.7 | 24.34 | 41124086.3 | 4.41684507 |
| T21 | 174671955 | 79.60 | 139056343.4 | 20.39 | 35615611.6 | 111.306157 |
| T22 | 119758503 | 80.60 | 96549305.12 | 19.38 | 23209197.9 | 13.7014124 |
| T23 | 161797790 | 79.20 | 128127669.9 | 20.81 | 33670120.1 | 0.65729026 |
| T25 | 168393125 | 77.00 | 129612188.3 | 23.03 | 38780936.7 | 2.1960895 |
| T30 | 191721412 | 78.90 | 151268194.1 | 21.10 | 40453217.9 | 4.07518997 |
| T31 | 182029273 | 79.70 | 144986315.9 | 20.35 | 37042957.1 | 22.6514933 |
| T32 | 120952287 | 79.50 | 96108687.25 | 20.54 | 24843599.7 | 8.00760456 |
| T33 | 154622501 | 78.70 | 121672446 | 21.31 | 32950055.0 | 34.637117 |
| Mean | **2233455546** | **78.67**  **(SE**$\boldsymbol{\pm0.40)}$ | **1784832731** | **21.33**  **(SE**$\boldsymbol{\pm0.40)}$ | **448622815** | **17.8218647** |

**Table S6. GenBank accession numbers and confirmed identities of viral sequences detected in *A. mellifera* and *B. terrestris* samples across Tasmanian sites. Viral identities were confirmed using BLASTn against the NCBI database, with sequences showing** $\geq$**90% identity considered true viral matches.**

| **Virus** | **Accession number** | **Pairwise identity (%)** |
| --- | --- | --- |
| Gorebridge virus | MH614301 | 97.10 |
| *Medicago sativa* Alphapartitivirus | PP333053 | 95.00 |
| Alfamovirus 1 | LC485018 | 98.60 |
| Aphid Lethal Paralysis virus | JX045858 | 95.50 |
| Victoria Bee virus | MG995723 | 96.70 |
| Ageratum Latent Ilarvirus | MT474112 | 98.60 |
| Lake Sinai virus 3 (LSV-3) | ON648746 | 95.80 |
| Lake Sinai virus 1 (LSV-1) | NC_074992 | 94.40 |
| Sacbrood virus (SBV) | KY465676 | 98.70 |
| Verbana Latent virus | PP502869 | 98.60 |
| *Solanum nigrum* Ilarvirus | BK061614 | 98.30 |
| Almendravirus Arboretum | OK491516 | 99.80 |
| White Clover Cryptic virus 2 | MH427306 | 99.80 |
| White Clover Mosaic virus | NC_003820 | 93.80 |
| Mentha Macluravirus | OL472141 | 91.50 |
| Moku virus | MF346349 | 98.80 |
| *Vespa velutina* Acypti-like virus | MN565042 | 97.20 |
| Boghill Burn virus | MH614292 | 98.10 |
| Hubei Picorna-like virus | KX883290 | 94.50 |
| Strawberry Latent Ringspot virus | MZ405640 | 95.60 |
| Red Clover Nepovirus A | NC_040400 | 92.70 |
| Arabis Mosaic virus DSMZ PV-0215 | OR477266 | 91.00 |
| Papua New Guinea Bee virus | MT482495 | 90.00 |
| Black Queen Cell virus (BQCV) | MF623171 | 94.40 |

**Table S7.** Summary of environmental vector fitting for NMDS ordinations of total, insect, and plant viral community composition in A. mellifera. Values represent p-values and correlation coefficients (r²) for the association between each environmental variable and NMDS ordination axes, based on 999 permutations. Statistically significant relationships (p $\leq$ 0.05) are shown in bold. Refer to Figs. 4 and S3 for corresponding NMDS ordination plots.

| *NMDS ORDINATION ENVFIT – TOTAL VIRAL COMMUNITY COMPOSITION* | | |
| --- | --- | --- |
| *Environmental factors* | *p* | *r^2^* |
| Mean annual temperature | 0.91 | 0.02 |
| Mean annual precipitation | 0.13 | 0.31 |
| Average summer wind velocity | 0.49 | 0.12 |
| Percentage of pasture | 0.16 | 0.29 |
| Longitude (X) | 0.93 | 0.01 |
| Latitude (Y) | 0.67 | 0.07 |
| *NMDS ORDINATION ENVFIT – PLANT VIRAL COMMUNITY COMPOSITION* | | |
| *Environmental factors* | *p* | *r^2^* |
| Mean annual temperature | 0.58 | 0.12 |
| **Mean annual precipitation** | **0.01** | **0.56** |
| Average summer wind velocity | 0.87 | 0.02 |
| Percentage of pasture | 0.65 | 0.06 |
| Longitude (X) | 0.39 | 0.15 |
| Latitude (Y) | 0.77 | 0.05 |
| *NMDS ORDINATION ENVFIT – INSECT VIRAL COMMUNITY COMPOSITION* | | |
| *Environmental factors* | *p* | *r^2^* |
| Mean annual temperature | 0.88 | 0.02 |
| **Mean annual precipitation** | **0.03** | **0.52** |
| Average summer wind velocity | 0.30 | 0.23 |
| Percentage of pasture | 0.51 | 0.12 |
| Longitude (X) | 0.22 | 0.23 |
| Latitude (Y) | 0.56 | 0.10 |

**Table S8.** Summary of environmental vector fitting for NMDS ordinations of total, insect, and plant viral community composition in B. terrestris. Values represent p-values and correlation coefficients (r²) for the association between each environmental variable and NMDS ordination axes, based on 999 permutations. Statistically significant relationships (p $\leq$ 0.05) are shown in bold. Refer to Figs. 4 and S3 for corresponding NMDS ordination plots.

| *NMDS ORDINATION ENVFIT – TOTAL VIRAL COMMUNITY COMPOSITION* | | |
| --- | --- | --- |
| *Environmental factors* | *p* | *r^2^* |
| Mean annual temperature | 0.25 | 0.24 |
| Mean annual precipitation | 0.23 | 0.25 |
| Average summer wind velocity | 0.42 | 0.16 |
| Percentage of pasture | 0.57 | 0.10 |
| Longitude (X) | 0.47 | 0.14 |
| Latitude (Y) | 0.38 | 0.18 |
| *NMDS ORDINATION ENVFIT – PLANT VIRAL COMMUNITY COMPOSITION* | | |
| *Environmental factors* | *p* | *r^2^* |
| Mean annual temperature | 0.81 | 0.05 |
| Mean annual precipitation | 0.09 | 0.44 |
| Average summer wind velocity | 0.63 | 0.11 |
| Percentage of pasture | 0.78 | 0.05 |
| Longitude (X) | 0.77 | 0.07 |
| Latitude (Y) | 0.75 | 0.75 |
| *NMDS ORDINATION ENVFIT – INSECT VIRAL COMMUNITY COMPOSITION* | | |
| *Environmental factors* | *p* | *r^2^* |
| Mean annual temperature | 0.92 | 0.02 |
| Mean annual precipitation | 0.15 | 0.33 |
| **Average summer wind velocity** | **0.04** | **0.43** |
| **Percentage of pasture** | **0.05** | **0.45** |
| **Longitude (X)** | **0.05** | **0.45** |
| Latitude (Y) | 0.06 | 0.43 |

**Table S9.** Correlations between alpha diversity (Shannon diversity and Chao1 richness) of the total, insect-associated, and plant-associated viromes in A. mellifera and environmental variables. Statistically significant relationships are highlighted in bold. Abbreviations: Temp = Mean annual temperature ($℃)$, Rain = Mean annual precipitation (mm), Wind = Average velocity of summer wind (m/s), Pasture = Percentage of pasture (%), X = Longitude, Y = Latitude.

| *Response variables* | *Predictor variables* | *p* | *R^2^* |
| --- | --- | --- | --- |
| Shannon diversity of total virome | Temp | 0.11 | 0.19 |
| Shannon diversity of total virome | Rain | 0.23 | 0.12 |
| Shannon diversity of total virome | Wind | 0.08 | 0.24 |
| Shannon diversity of total virome | Pasture | 0.86 | 0.003 |
| Shannon diversity of total virome | X | 0.24 | 0.11 |
| Shannon diversity of total virome | Y | 0.78 | 0.01 |
| Chao1 richness of total virome | Temp | 0.08 | 0.23 |
| Chao1 richness of total virome | Rain | 0.27 | 0.09 |
| Chao1 richness of total virome | Wind | 0.52 | 0.04 |
| Chao1 richness of total virome | Pasture | 0.95 | 0.003 |
| Chao1 richness of total virome | X | 0.16 | 0.16 |
| Chao1 richness of total virome | Y | 0.61 | 0.02 |
| Shannon diversity of insect virome | Temp | 0.85 | 0.003 |
| **Shannon diversity of insect virome** | **Rain** | **0.05** | **0.30** |
| Shannon diversity of insect virome | Wind | 0.47 | 0.04 |
| Shannon diversity of insect virome | Pasture | 0.4 | 0.06 |
| Shannon diversity of insect virome | X | 0.99 | 3.61^8^ |
| Shannon diversity of insect virome | Y | 0.25 | 0.11 |
| Chao1 richness of insect virome | Temp | 0.51 | 0.04 |
| Chao1 richness of insect virome | Rain | 0.52 | 0.03 |
| Chao1 richness of insect virome | Wind | 0.31 | 0.09 |
| Chao1 richness of insect virome | Pasture | 0.32 | 0.08 |
| Chao1 richness of insect virome | X | 0.5 | 0.04 |
| Chao1 richness of insect virome | Y | 0.15 | 0.16 |
| Shannon diversity of plant virome | Temp | 0.47 | 0.05 |
| Shannon diversity of plant virome | Rain | 0.12 | 0.19 |
| Shannon diversity of plant virome | Wind | 0.65 | 0.02 |
| Shannon diversity of plant virome | Pasture | 0.35 | 0.07 |
| Shannon diversity of plant virome | X | 0.11 | 0.20 |
| Shannon diversity of plant virome | Y | 0.89 | 0.001 |
| **Chao1 richness of plant virome** | **Temp** | **0.05** | **0.28** |
| **Chao1 richness of plant virome** | **Rain** | **0.001** | **0.64** |
| Chao1 richness of plant virome | Wind | 0.71 | 0.01 |
| Chao1 richness of plant virome | Pasture | 0.11 | 0.20 |
| Chao1 richness of plant virome | X | 0.17 | 0.15 |
| Chao1 richness of plant virome | Y | 0.26 | 0.10 |

**Table S10.** Correlations between alpha diversity (Shannon diversity and Chao1 richness) of the total, insect-associated, and plant-associated viromes in B. terrestris and environmental variables. Statistically significant relationships are highlighted in bold. Abbreviations: Temp = Mean annual temperature ($℃)$, Rain = Mean annual precipitation (mm), Wind = Average velocity of summer wind (m/s), Pasture = Percentage of pasture (%), X = Longitude, Y = Latitude.

| *Response variables* | *Predictor variables* | *p* | *R^2^* |
| --- | --- | --- | --- |
| Shannon diversity of total virome | Temp | 0.99 | 2.37^-6^ |
| **Shannon diversity of total virome** | **Rain** | **0.05** | **0.27** |
| Shannon diversity of total virome | Wind | 0.75 | 0.01 |
| Shannon diversity of total virome | Pasture | 0.86 | 0.003 |
| Shannon diversity of total virome | X | 0.39 | 0.07 |
| Shannon diversity of total virome | Y | 0.44 | 0.05 |
| Chao1 richness of total virome | Temp | 0.88 | 0.07 |
| **Chao1 richness of total virome** | **Rain** | **0.01** | **0.45** |
| Chao1 richness of total virome | Wind | 0.54 | 0.04 |
| **Chao1 richness of total virome** | **Pasture** | **0.01** | **0.48** |
| Chao1 richness of total virome | X | 0.43 | 0.06 |
| Chao1 richness of total virome | Y | 0.51 | 0.04 |
| Shannon diversity of insect virome | Temp | 0.09 | 0.23 |
| Shannon diversity of insect virome | Rain | 0.15 | 0.18 |
| Shannon diversity of insect virome | Wind | 0.86 | 0.003 |
| Shannon diversity of insect virome | Pasture | 0.13 | 0.2 |
| Shannon diversity of insect virome | X | 0.39 | 0.07 |
| Shannon diversity of insect virome | Y | 0.61 | 0.03 |
| Chao1 richness of insect virome | Temp | 0.88 | 0.002 |
| **Chao1 richness of insect virome** | **Rain** | **0.03** | **0.38** |
| Chao1 richness of insect virome | Wind | 0.34 | 0.08 |
| Chao1 richness of insect virome | Pasture | 0.16 | 0.17 |
| **Chao1 richness of insect virome** | **X** | **0.04** | **0.31** |
| Chao1 richness of insect virome | Y | 0.55 | 0.03 |
| **Shannon diversity of plant virome** | **Temp** | **0.05** | **0.3** |
| **Shannon diversity of plant virome** | **Rain** | **0.05** | **0.28** |
| Shannon diversity of plant virome | Wind | 0.81 | 0.005 |
| Shannon diversity of plant virome | Pasture | 0.08 | 0.25 |
| Shannon diversity of plant virome | X | 0.89 | 0.001 |
| Shannon diversity of plant virome | Y | 0.15 | 0.18 |
| Chao1 richness of plant virome | Temp | 0.24 | 0.12 |
| Chao1 richness of plant virome | Rain | 0.08 | 0.25 |
| Chao1 richness of plant virome | Wind | 0.83 | 0.002 |
| **Chao1 richness of plant virome** | **Pasture** | **0.01** | **0.48** |
| Chao1 richness of plant virome | X | 0.74 | 0.01 |
| Chao1 richness of plant virome | Y | 0.11 | 0.21 |

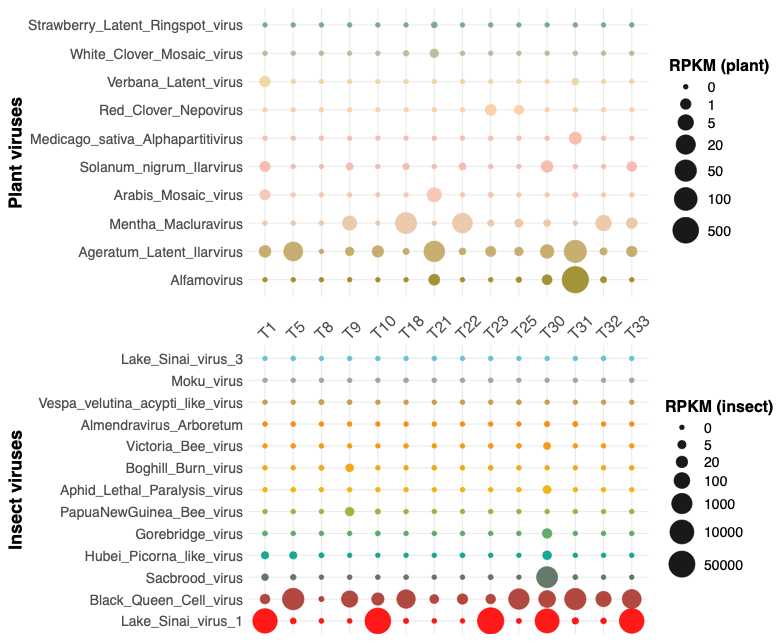

**Figure S1.** Bubble plot showing the RPKM (Reads Per Kilobase per Million mapped reads) normalised abundance of plant and insect viruses detected across Tasmanian A. mellifera samples. Each row corresponds to an individual virus (top panel: plant viruses; bottom panel: insect viruses), while each column represents a sampling site along the shared x-axis. Bubble size is proportional to the RPKM value for each virus at each site, with larger bubbles indicating higher abundance. Legends to the right denote the RPKM scales used for each virus group, with ranges adjusted to reflect the differing abundance levels of plant versus insect viruses.

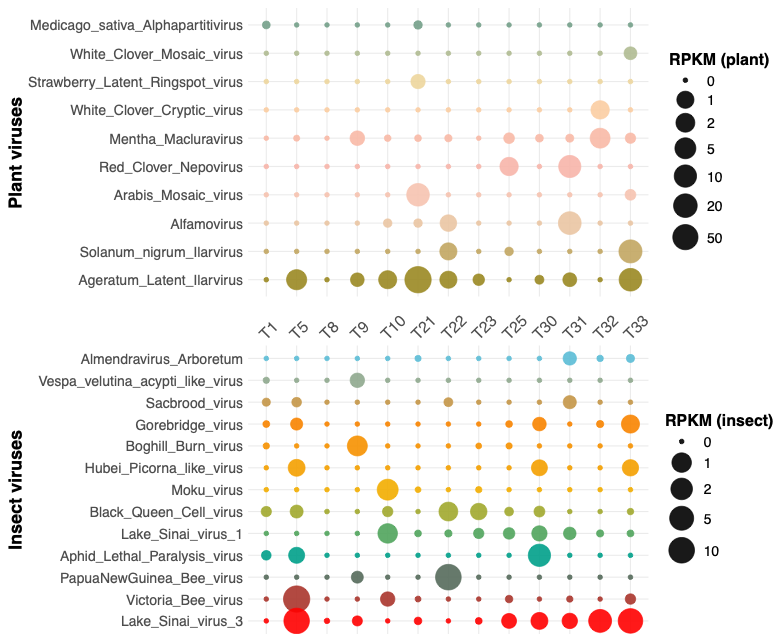

**Figure S2.** Bubble plot showing the RPKM (Reads Per Kilobase per Million mapped reads) normalised abundance of plant and insect viruses detected across Tasmanian B. terrestris samples. Each row corresponds to an individual virus (top panel: plant viruses; bottom panel: insect viruses), while each column represents a sampling site along the shared x-axis. Bubble size is proportional to the RPKM value for each virus at each site, with larger bubbles indicating higher abundance. Legends to the right denote the RPKM scales used for each virus group, with ranges adjusted to reflect the differing abundance levels of plant versus insect viruses.

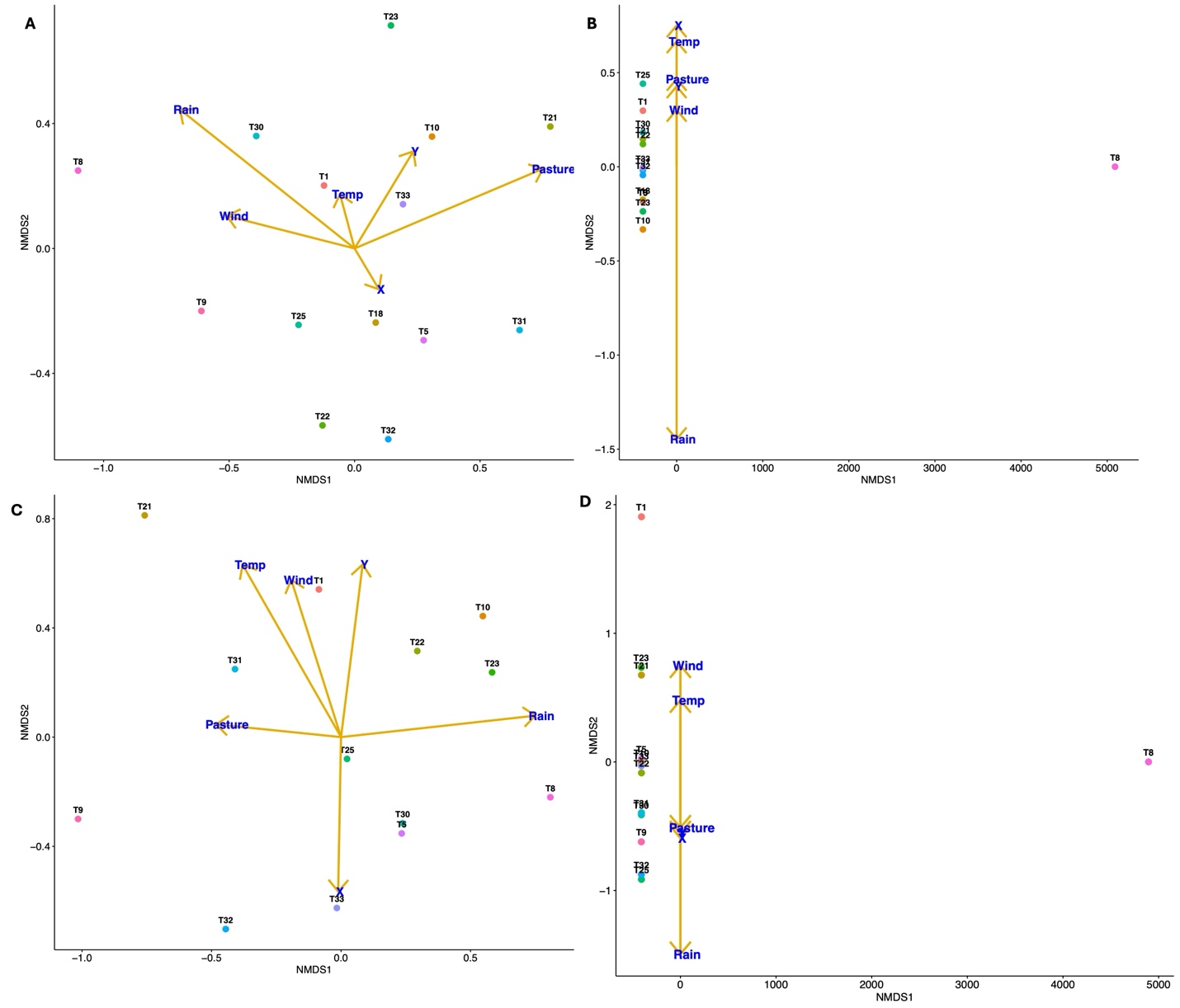

**Figure S3.** NMDS ordinations of viral community composition across Tasmanian sites based on Bray–Curtis dissimilarities in (A) A. mellifera total virome (Stress = 0.19), (B) A. mellifera plant-associated virome (Stress = 0.0002), (C) B. terrestris total virome (Stress = ), and (D) B. terrestris plant-associated virome (Stress = …). Environmental vector correlations (envfit) are provided in Tables S6 and S7 for A. mellifera and B. terrestris, respectively. Abbreviations: Temp = mean annual temperature (°C), Rain = mean annual precipitation (mm), Wind = average summer wind velocity (m/s), Pasture = percentage of pasture (%), X = longitude, Y = latitude. See Fig. 4 for NMDS ordinations of insect virome compositions for A. mellifera and B. terrestris, and Tables S7 and S8 for the corresponding envfit correlations.

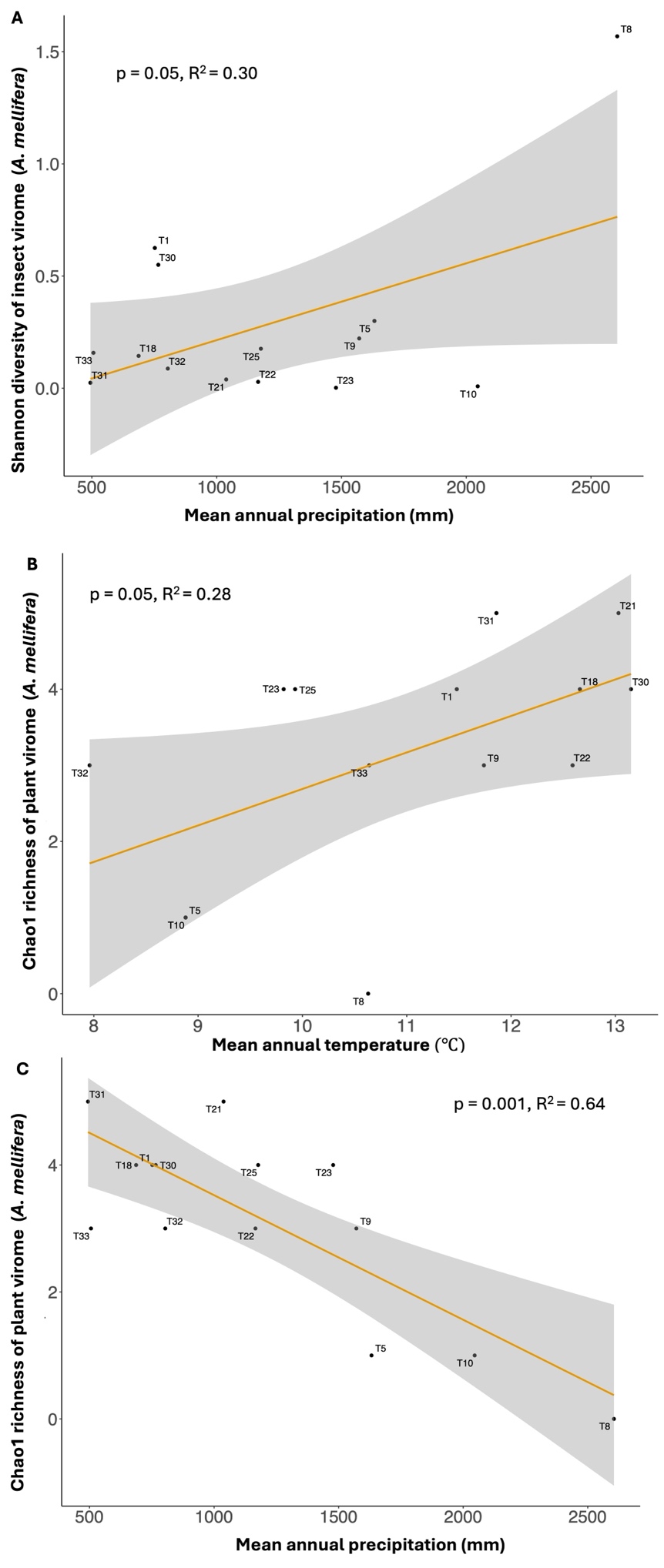

**Figure S4.** Linear relationships in A. mellifera showing: (A) a positive correlation between insect viral diversity and mean annual precipitation, (B) a positive correlation between plant viral richness and mean annual temperature, and (C) a negative correlation between plant viral richness and mean annual precipitation. Diversity measure = Shannon; Richness measure = Chao1.

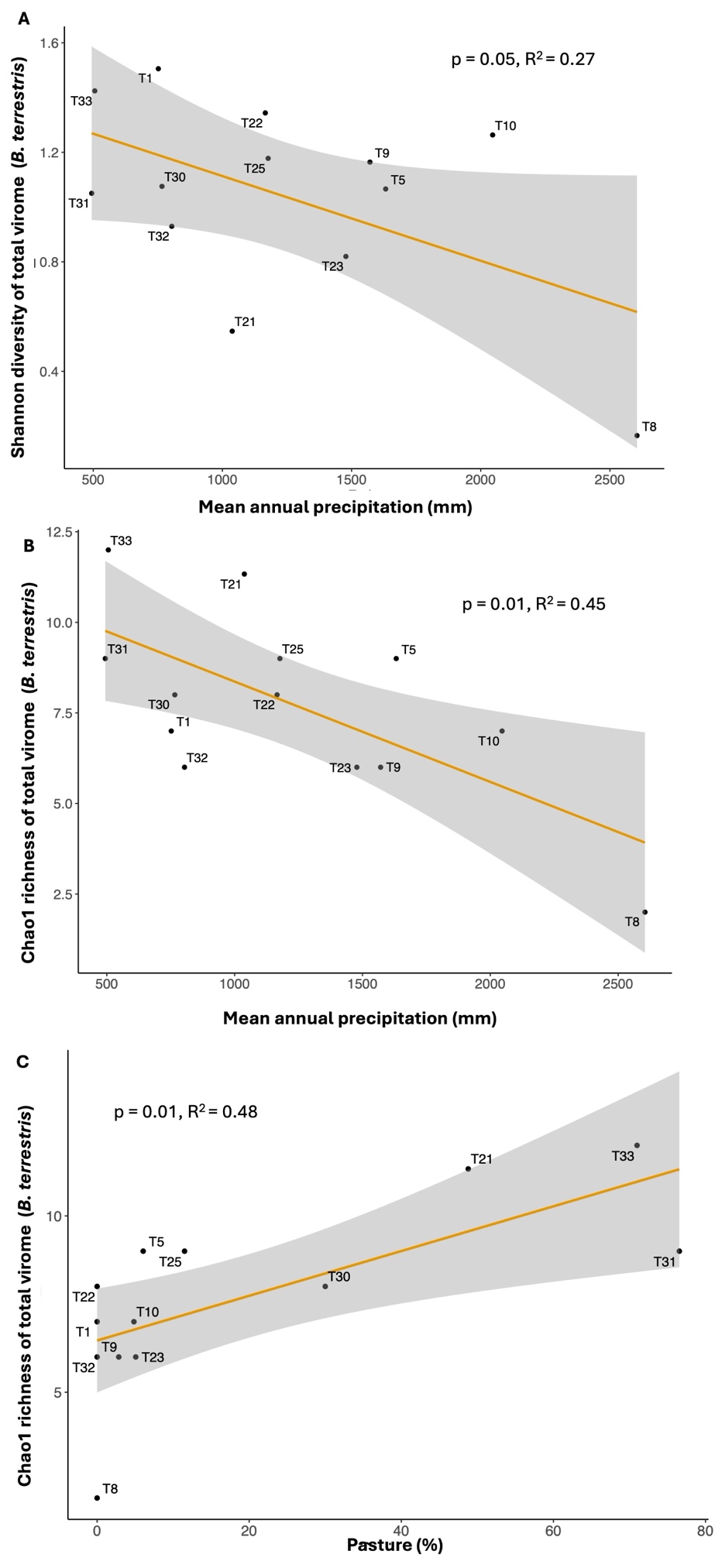

**Figure S5.** Linear relationships in *B. terrestris* showing: (A) a negative correlation between total viral diversity and mean annual precipitation, (B) a negative correlation between total viral richness and mean annual precipitation, and (C) a positive correlation between total viral richness and percentage of pasture. Diversity measure = Shannon; Richness measure = Chao1.

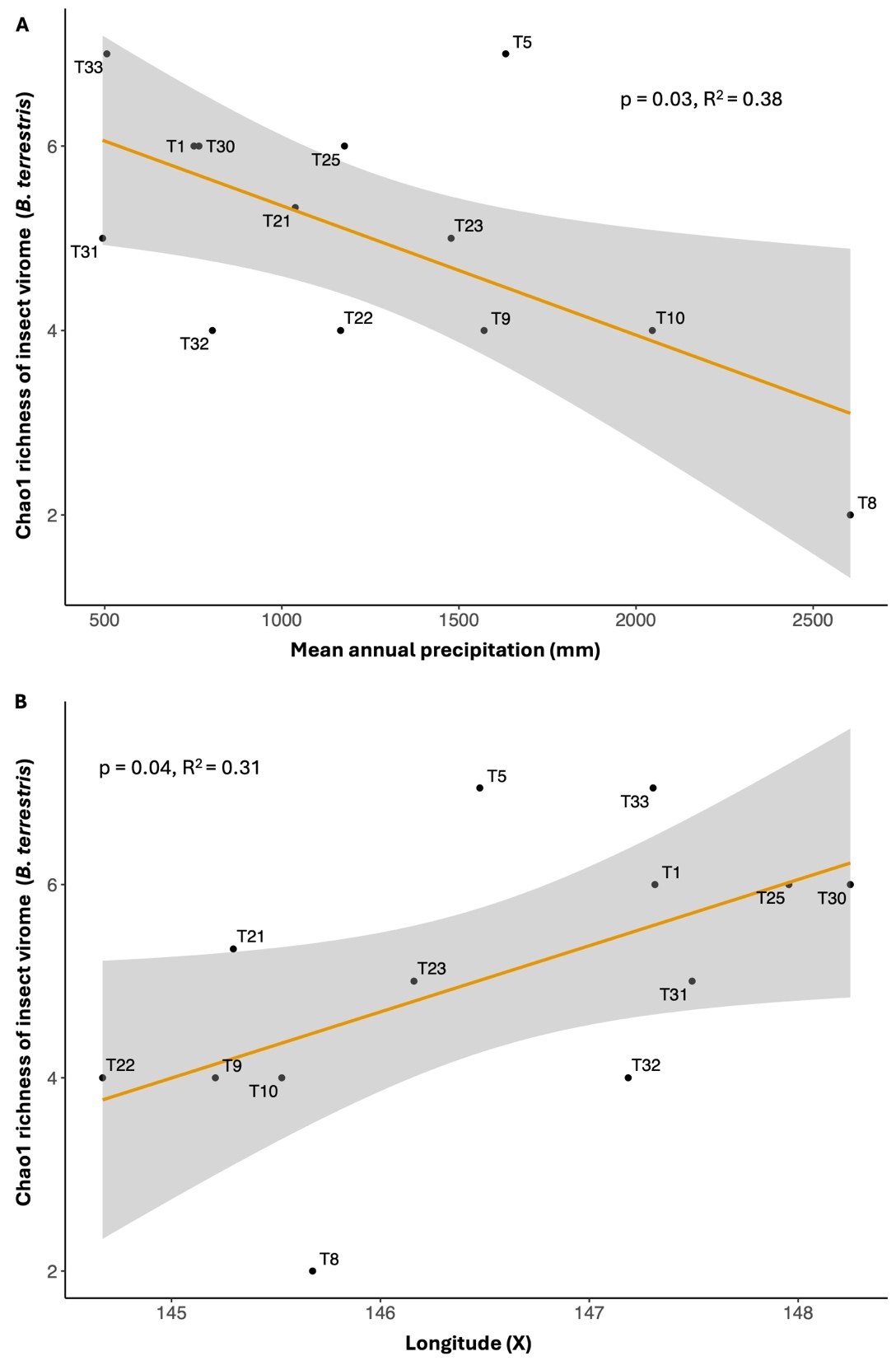

**Figure S6.** Linear relationships in *B. terrestris* showing: (A) a negative correlation between insect viral richness and mean annual precipitation, (B) a positive correlation between insect viral richness and longitude. Richness measure = Chao1.

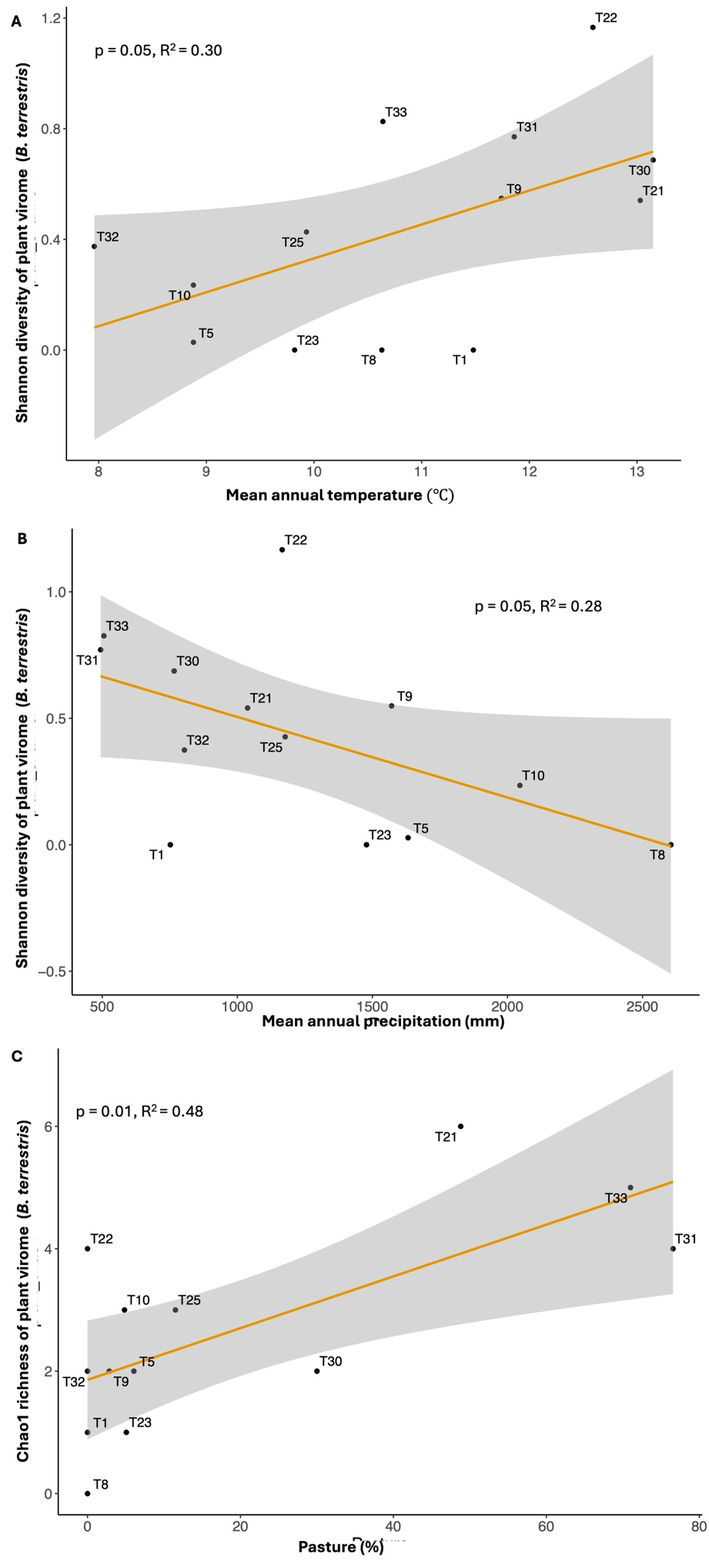

**Figure S7.** Linear relationships in *B. terrestris* showing: (A) a positive correlation between plant viral diversity and mean annual temperature, (B) a negative correlation between plant viral diversity and mean annual precipitation, (C) a positive correlation between plant viral richness and pasture percentage. Diversity measure = Shannon; Richness measure = Chao1.

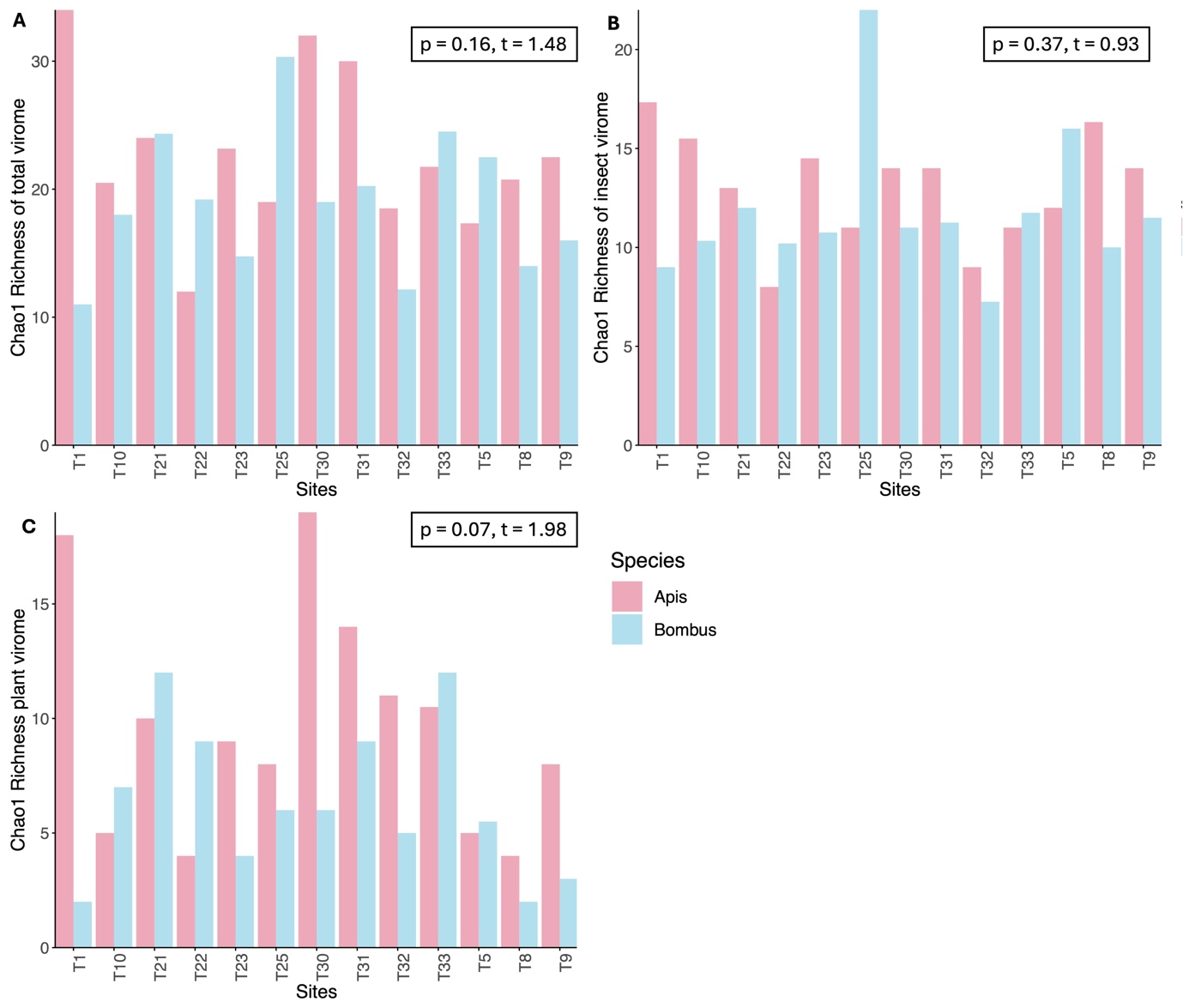
**Figure S8.** Comparison of Chao1 richness of viruses between *A. mellifera* and *B. terrestris* across sites: (A) total virome richness, (B) insect virome richness and (C) plant virome richness. Paired t-tests are shown in each panel, with no significant differences observed between species (p > 0.05).

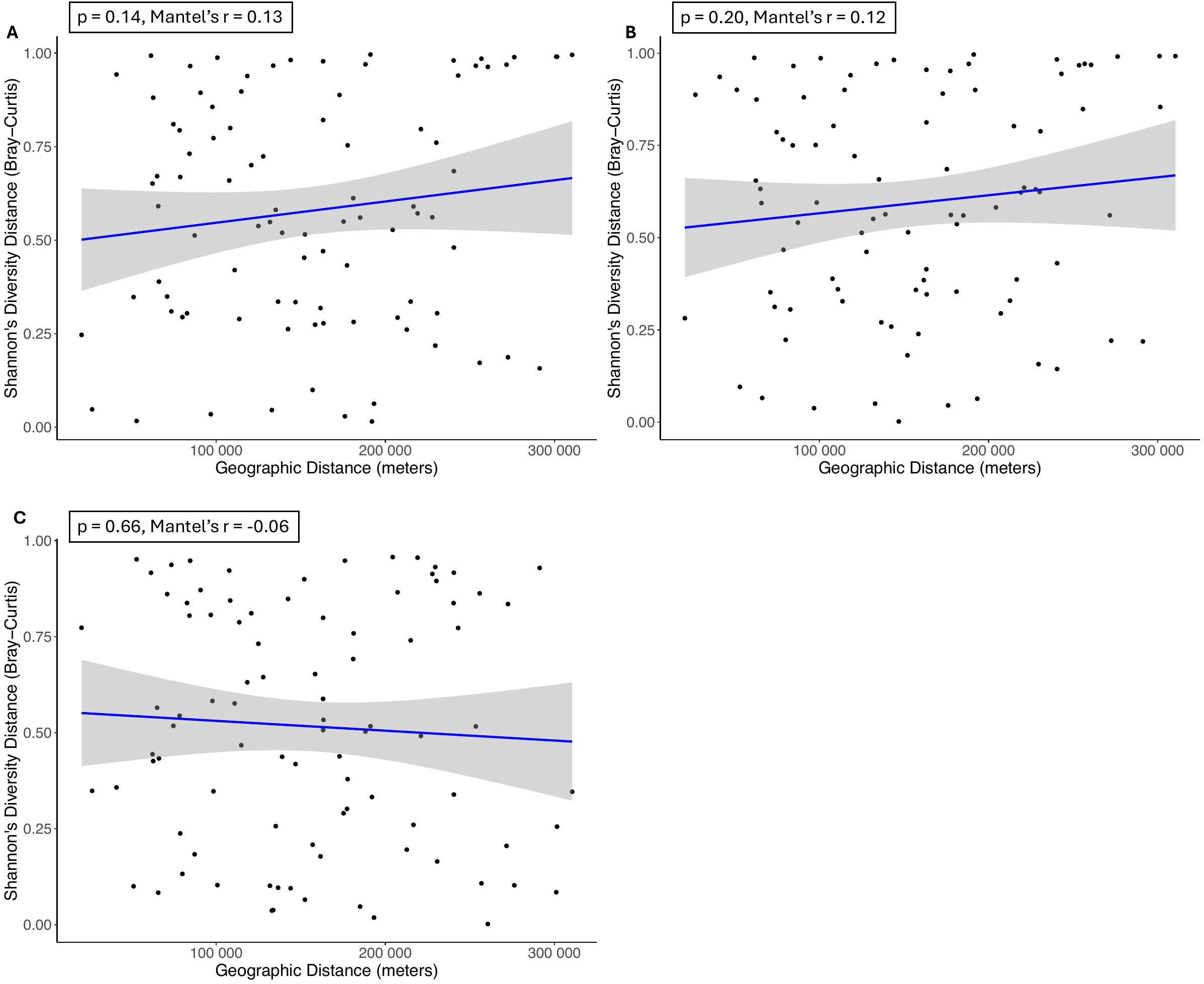

**Figure S9.** Tests of isolation by distance for *A. mellifera* virome diversity across sites. Mantel tests illustrate the relationship between geographic distance (in meters) and pairwise differences (using Bray–Curtis) in Shannon’s diversity for (A) total virome, (B) insect virome and (C) plant virome. Each point represents a pairwise comparison between sites; shaded areas represent 95% confidence intervals with the blue regression line. No significant relationships were observed (p > 0.05).

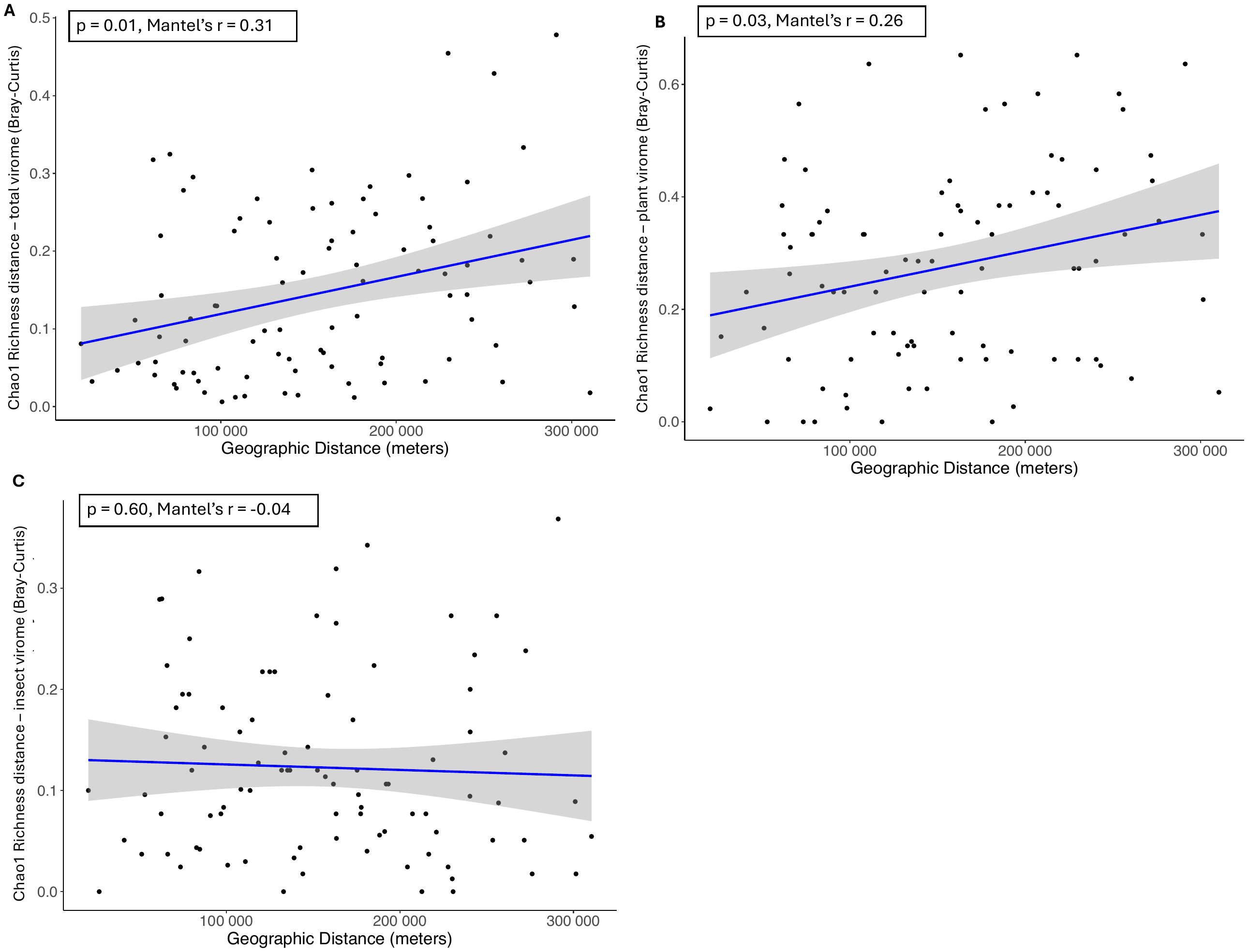

**Figure S10.** Isolation by distance analysis for *A. mellifera* virome richness across sites. Mantel tests illustrate the relationship between geographic distance (in meters) and pairwise differences (using Bray–Curtis) in Chao1 richness for (A) total virome, (B) plant virome, and (C) insect virome. Each point represents a pairwise comparison between sites; shaded areas represent 95% confidence intervals with the blue regression line. Significant positive relationship was observed in (A) and (B) for total and plant viromes (p $\leq$0.05), but not in (C) for insect virome (p > 0.05).

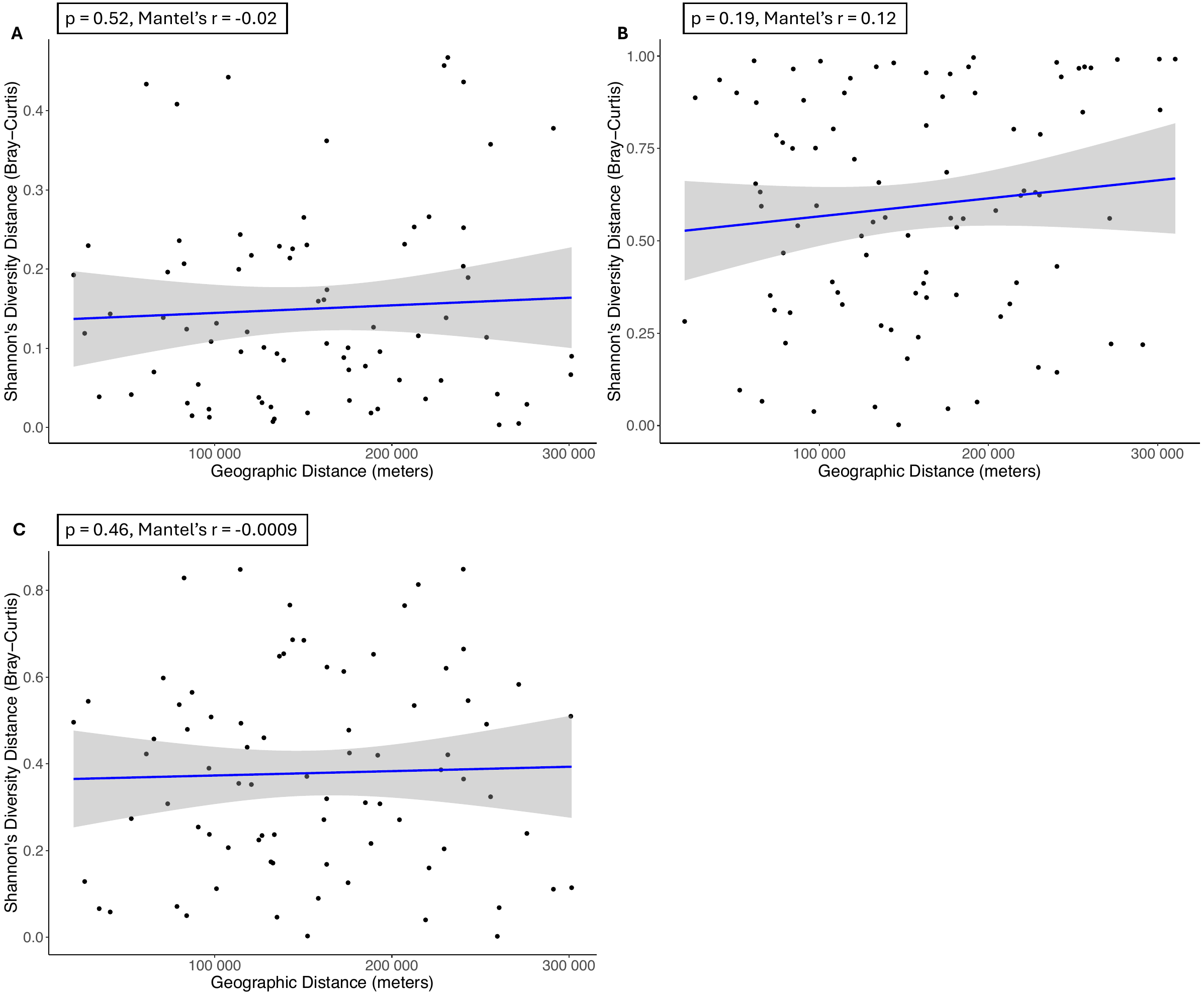
 **Figure S11.** Tests of isolation by distance for *B. terrestris* virome diversity across sites. Mantel tests illustrate the relationship between geographic distance (in meters) and pairwise differences (using Bray–Curtis) in Shannon’s diversity for (A) total virome, (B) insect virome and (C) plant virome. Each point represents a pairwise comparison between sites; shaded areas represent 95% confidence intervals with the blue regression line. No significant relationship was observed (p > 0.05).

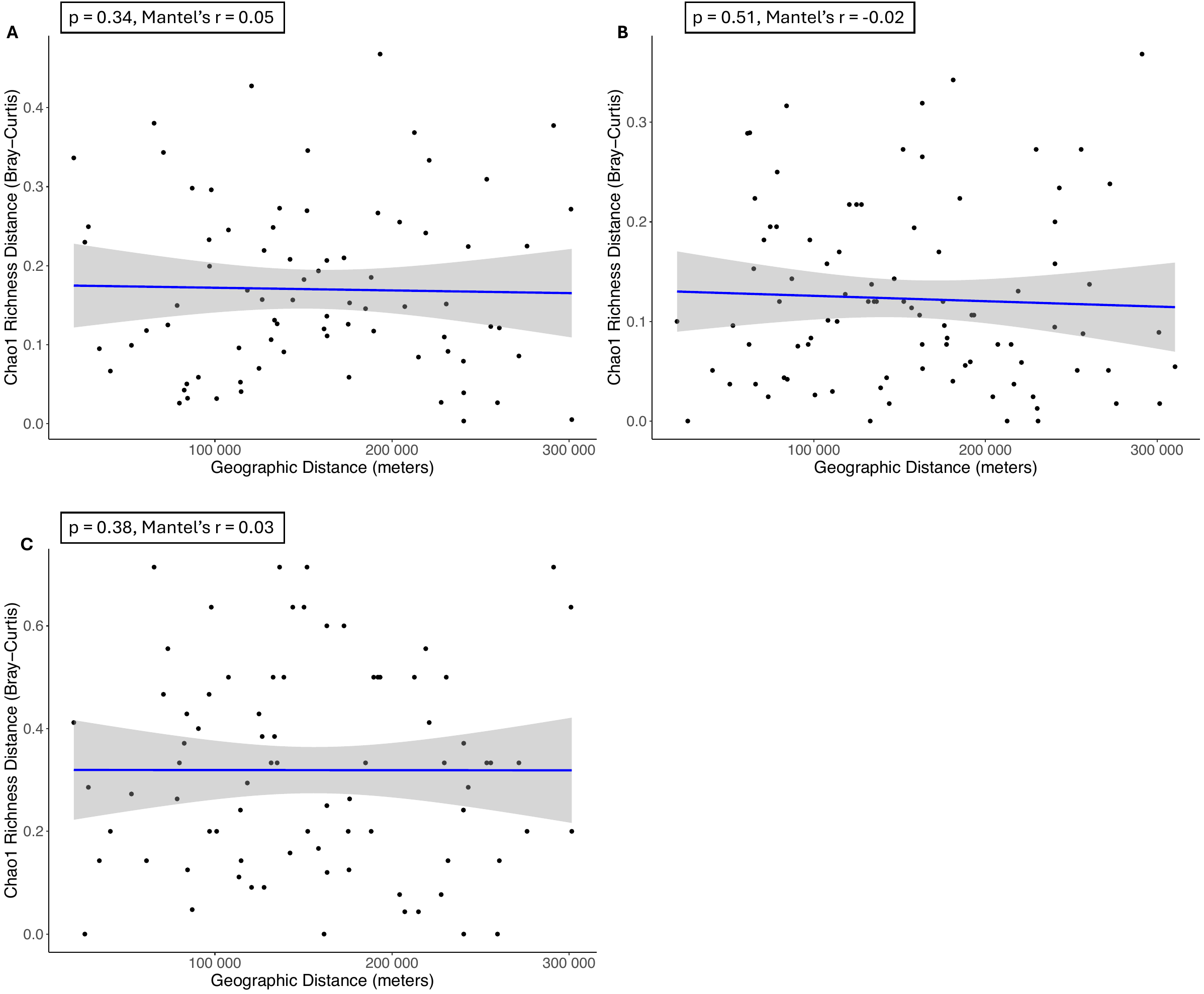

**Figure S12.** Tests of isolation by distance for *B. terrestris* virome richness across sites. Mantel tests illustrate the relationship between geographic distance (in meters) and pairwise differences (using Bray–Curtis) in Chao1 richness for (A) total virome, (B) insect virome and (C) plant virome. Each point represents a pairwise comparison between sites; shaded areas represent 95% confidence intervals with the blue regression line. No significant relationship was observed (p > 0.05).

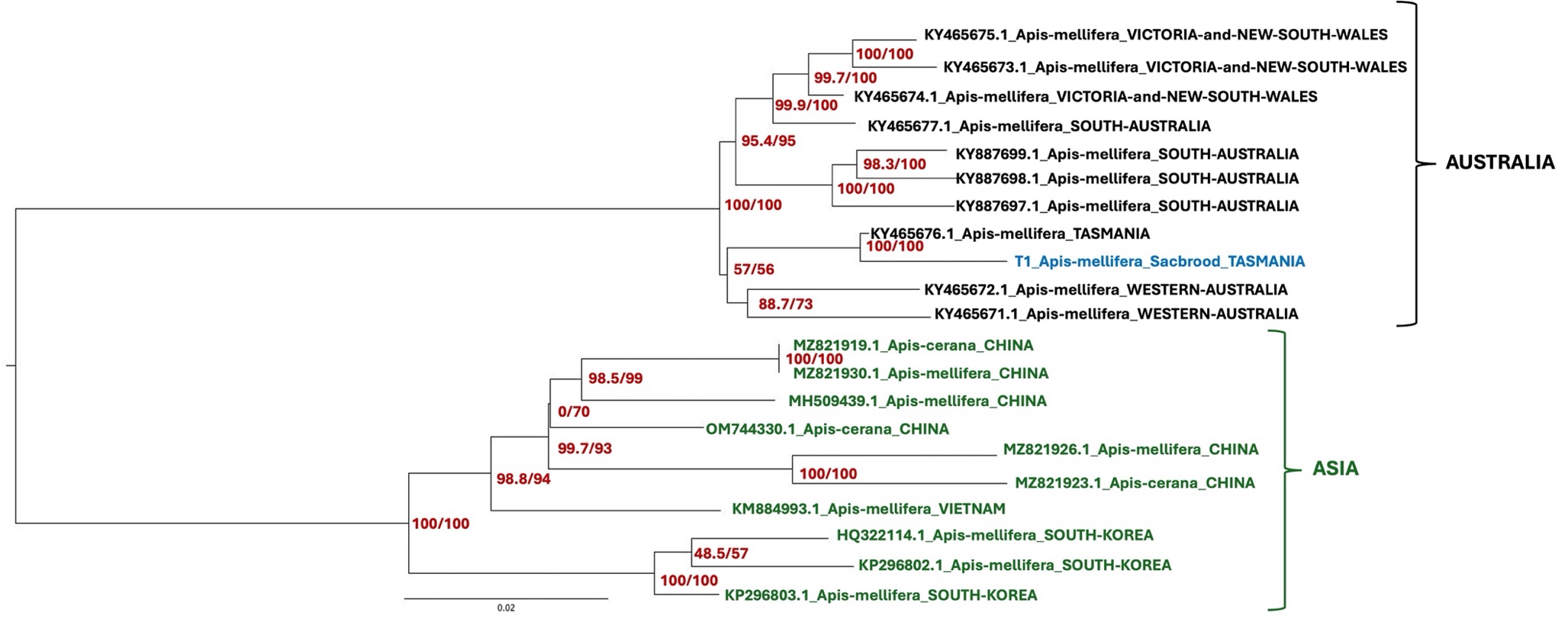

**Figure S13.** Sacbrood virus (SBV) phylogenetic tree. The phylogenetic tree was generated using maximum likelihood in IQ-TREE (Trifinopoulos et al., 2016), with the GTR+F+G4 substitution model which had the optimal BIC score, as determined by ModelFinder (Kalyaanamoorthy et al., 2017). Branch supports were estimated using Ultrafast bootstrap approximation (UFBoot; Hoang et al., 2017) using 1000 replicates. Support values shown are SH-like approximate likelihood ratio test (SH-aLRT) and UFBoot supports.
